# An automated high-content synaptic phenotyping platform in human neurons and astrocytes reveals a role for BET proteins in synapse assembly

**DOI:** 10.1101/2022.05.17.492322

**Authors:** Martin H. Berryer, Gizem Rizki, Sara G. Susco, Daisy Lam, Angelica Messana, Darina Trendafilova, Anna Nathanson, Kyle W. Karhohs, Beth A. Cimini, Kathleen Pfaff, Anne E. Carpenter, Lee L. Rubin, Lindy E. Barrett

## Abstract

Resolving fundamental molecular and functional processes underlying human synaptic development is crucial for understanding normal brain function as well as dysfunction in disease. Based upon increasing evidence of species divergent features of brain cell types, coupled with emerging studies of complex human disease genetics, we developed the first automated and quantitative high-content synaptic phenotyping platform using human neurons and astrocytes. To establish the robustness of our platform, we screened the effects of 376 small molecules on presynaptic density, neurite outgrowth and cell viability, validating six small molecules which specifically enhanced human presynaptic density *in vitro*. Astrocytes were essential for mediating the effects of all six small molecules, underscoring the relevance of non-cell autonomous factors in synapse assembly and their importance in synaptic screening applications. Bromodomain and extraterminal (BET) inhibitors emerged as the most prominent hit class and global transcriptional analyses using multiple BET inhibitors confirmed upregulation of synaptic gene expression programs. Through these analyses, we demonstrate the robustness of our automated screening platform for identifying potent synaptic modulators, which can be further leveraged for scaled analyses of human synaptic mechanisms and drug discovery efforts.

## INTRODUCTION

Synapses are inter-cellular junctions between neurons crucial for information processing. Seminal studies using the frog neuromuscular junction (Del Castillo and Katz, 1954) or giant squid synapse (Bullock and Hagiwara, 1957) established fundamental principles of chemical transmission. Subsequent genetic and neurobiological studies have further illuminated our understanding of the mammalian central nervous system and have implicated synaptic alterations in brain disorders including autism spectrum disorder (Bourgeron, 2015; De Rubeis et al., 2014), schizophrenia (Fromer et al., 2014; Kirov et al., 2012; Purcell et al., 2014), and Alzheimer’s disease (Kunkle et al., 2019; Spires-Jones and Hyman, 2014). Thus, insights into synaptic development and function are critical for our understanding of normal brain function as well as dysfunction in disease. Diverse genes such as neuroligins and neurexins (Graf et al., 2004; Varoqueaux et al., 2006), neurotrophic receptors (Elmariah et al., 2004), EphrinB and EphrinB receptors (Henderson and Dalva, 2018) and cadherins, protocadherins and catenins (Arikkath and Reichardt, 2008; Weiner et al., 2005; Yamagata et al., 2018) have been implicated in synaptic biology, although given the complex nature of mammalian brain development and function, molecular programs underlying neuronal connectivity have yet to be fully resolved.

Notably, most synaptic studies in mammals use rodent primary neuronal cultures (Linhoff et al., 2009; Nieland et al., 2014; Sharma et al., 2013; Sharma et al., 2012; Verstraelen et al., 2014). While the general principles of neuronal and synaptic development are thought to be conserved across species (Defelipe, 2011), increasing evidence suggests that species-divergent features influence signal processing in dendrites and axons which in turn may contribute to differences in cognition and function across organisms. Indeed, it is possible that human-specific aspects of synapse biology contribute to higher order cognition and function and may be perturbed in disease, underscoring the importance of utilizing both highly tractable, well-studied animal models as well as emerging human cellular *in vitro* models to identify and analyze convergent and divergent principles. In this regard, studies comparing human and rodent cortical neurons have revealed differences in membrane capacitance (Eyal et al., 2016), epigenetic signatures (Luo et al., 2017), gene expression and transcriptional regulatory mechanisms (Bakken et al., 2016; Hodge et al., 2019; Zeng et al., 2012), as well as changes in activity-dependent gene expression (Pruunsild et al., 2017) associated with distinct somatic, axonal, dendritic and spine morphologies (Beaulieu-Laroche et al., 2018; Benavides-Piccione et al., 2002; Mohan et al., 2015) which correlate with differences in synaptic plasticity (Szegedi et al., 2016; Verhoog et al., 2013) and likely contribute to the differences observed in the integrative properties of the neuron (Molnar et al., 2016; Szegedi *et al*., 2016; Verhoog *et al*., 2013). For example, electrophysiological analyses have revealed key differences in spike timing-dependent plasticity in the human cortex compared with rodent, including the ability of human synapses to change strength bidirectionally in response to spike timing (Verhoog *et al*., 2013). A study of connections between human pyramidal neurons and fast-spiking GABAergic interneurons reported four times more functional pre-synaptic release sites in humans compared with rodents (Molnar *et al*., 2016). Paralleling functional differences, dendritic spines from pyramidal neurons in the human brain were reported to be larger, longer and more densely packed compared with their rodent counterparts (Benavides-Piccione *et al*., 2002). Moreover, astrocytes are known to play critical roles in synapse development, maintenance and function (Allen et al., 2012; Allen and Eroglu, 2017; Bernardinelli et al., 2014; Di Castro et al., 2011; Krencik et al., 2017; Panatier et al., 2011; Ullian et al., 2001), and human astrocytes exhibit transcriptomic profiles as well as functional and structural properties which diverge from rodent astrocytes (Hawrylycz et al., 2012; Oberheim et al., 2009). Thus, human and rodent astrocytes likely differ in their contributions to synaptic and neuronal function (Oberheim *et al*., 2009; Zhang et al., 2016). The above species-specific features highlight the importance of studying human neurons and human astrocytes to decode human nervous system function, which critically depends on networked neural activity involving both cell types.

Recent advances in human pluripotent stem cell (hPSC) biology and differentiation now allow for the generation of multiple human brain cell types *in vitro* including glutamatergic neurons (Busskamp et al., 2014; Chambers et al., 2009; Zhang et al., 2013). This has provided key insights into molecular mechanisms underlying normal synaptic development (Patzke et al., 2019) as well as alterations in human neurodevelopmental and neurological disorders such as autism (Krey et al., 2013; Marchetto et al., 2017; Yi et al., 2016), schizophrenia (Pak et al., 2015; Yoon et al., 2014), Alzheimer’s disease (Israel et al., 2012; Lin et al., 2010) and Parkinson’s disease (Sanchez-Danes et al., 2012; Vera et al., 2016). Advances in differentiation technologies have also facilitated the scalable production of hPSC-derived neurons to allow for phenotypic cell-based screening assays. Both genetic and pharmacological drug screening strategies using hPSC derived neurons have been employed to study pathways regulating Tau and β-amyloid production in Alzheimer’s disease (Brownjohn et al., 2017; Kondo et al., 2017; van der Kant et al., 2019; Wang et al., 2017). Other studies have used hPSC-derived neurons to measure neurotoxicity (Ryan et al., 2016), neurite outgrowth (Sherman and Bang, 2018) and gene function using CRISPR interference and activation (Tian et al., 2021; Tian et al., 2019), further supporting the utility of such models for high-throughput screens. However, despite their unprecedented potential, scaled analyses of synaptic mechanisms using human cellular systems have not yet been achieved, in part due to the technical challenges associated with generating and analyzing reproducible culture preparations essential for these analyses. Indeed, while studies using human neurons have quantified the effects of candidate small molecules on synaptic density (Green et al., 2019; Patzke *et al*., 2019; Trujillo et al., 2021), to date we are not aware of any automated systems for their unbiased interrogation. This will become increasingly relevant for the study of basic synaptic mechanisms as additional species divergent features of brain cell types are identified, and for the study of complex human diseases which impact synaptic function.

We therefore devised a fully automated and quantitative high-content synaptic phenotyping platform using *in vitro* derived human glutamatergic neurons and primary human astrocytes. To demonstrate the utility of our platform, we screened effects of 376 small molecules on presynaptic density, neurite outgrowth and cell viability. These analyses identified six small molecules which specifically enhanced presynaptic density. We then probed the most prominent hit class, bromodomain and extraterminal (BET) inhibitors, through global transcriptional analyses. While BET inhibitors have been associated with decreased synaptic gene expression in rodent models, we confirmed upregulation of synaptic gene expression programs in human cellular models, supporting the results of our synaptic assay. Taken together, we have established a novel human synaptic phenotyping platform which effectively identifies synaptic modulators *in vitro*. This platform can now be leveraged for further interrogation of human synaptic mechanisms and drug discovery efforts.

## RESULTS

### Generation of robust human neuron and astrocyte co-cultures for synaptic screening applications

To reproducibly generate large batches of human induced neurons (hNs) from hPSCs, we used a well-described *in vitro* differentiation protocol based on ectopic expression of Neurogenin 2 (Ngn2) combined with developmental patterning using small molecules (Nehme et al., 2018; Zhang *et al*., 2013). We selected this protocol based on extensive molecular and physiological characterization of these hNs alongside their use in multiple studies of disease-associated genes (Busskamp *et al*., 2014; Deneault et al., 2018; Lin et al., 2018; Meijer et al., 2019; Nehme *et al*., 2018; Pak *et al*., 2015; Yi *et al*., 2016; Zhang *et al*., 2013; Zhang et al., 2018) and in the only published genome-wide CRISPRi/CRISPRa screens of *in vitro* derived human neurons to date (Tian *et al*., 2021; Tian *et al*., 2019). Importantly, studies using these hNs have identified physiological and/or morphological phenotypes upon perturbation of synaptic genes (e.g., *NRXN1*, *SHANK3)*, which have then been recapitulated using other differentiation paradigms, further validating the accuracy of synaptic phenotypes detected in hNs generated with this protocol (Pak *et al*., 2015; Yi *et al*., 2016). Moreover, most if not all neuronal differentiation paradigms *in vitro* generate heterogeneous cell types at lower throughput (Chambers *et al*., 2009); however this protocol produces glutamatergic neurons at large scale that are highly homogeneous in terms of cell type, making it uniquely suited for screening applications (Nehme *et al*., 2018; Tian *et al*., 2021; Tian *et al*., 2019). Data from our laboratory and others confirm that hNs are electrophysiologically active by day 21 *in vitro* displaying spontaneous excitatory post-synaptic currents (sEPSCs) and NMDAR-mediated currents (Meijer *et al*., 2019; Nehme *et al*., 2018; Susco et al., 2020).

To overcome the limitations of lentiviral titer used to transduce Ngn2, and to enhance differentiation efficiency, we used a pair of TALENs to stably introduce a doxycycline inducible Ngn2 cassette (Zhang *et al*., 2013) into the AAVS1 safe harbor locus of the hPSC line H1 to generate a zeocin resistant doxycycline inducible Ngn2 hPSC line, referred to as iNgn2-hPSC (**Fig. 1a**), similar to the approach used by Meijer et al (Meijer *et al*., 2019). Upon transfection, cells were selected for geneticin resistance and individual clones were isolated, expanded, re-plated for genomic DNA extraction and PCR analysis of the transgene integration, and further analyzed by G-band karyotyping (**Supplemental Fig. 1a, b**). To generate large-scale neuronal preparations essential for screening applications, we then differentiated the iNgn2-hPSCs into hNs as described (Nehme *et al*., 2018) (**Fig. 1b**).

**Figure 1:**
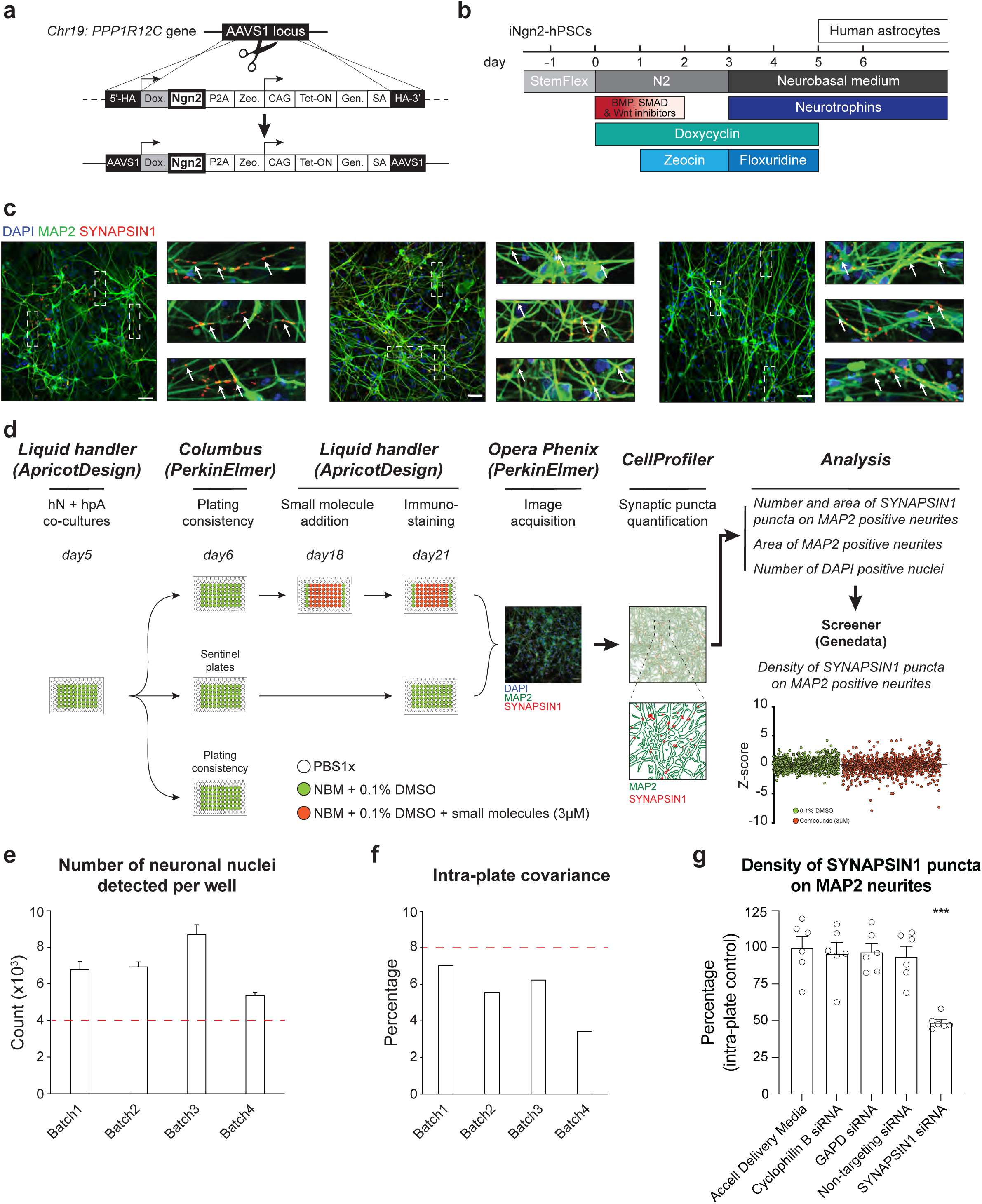
Development of an automated, high-content human synaptic phenotyping platform. **a.** Schematic of stable integration of a doxycycline responsive Ngn2 (iNgn2) cassette into hPSCs. TALENs were used to insert iNgn2 into the AAVS1 safe harbor locus of the *PPP1R12C* gene. **b.** Summary of human induced neuron (hN) protocol. Human pluripotent stem cells (hPSCs) are differentiated into neural progenitor like cells using extrinsic inhibition of BMP, SMAD and Wnt in combination with doxycycline driven transient ectopic Ngn2 expression. Zeocin is used as a selective agent. Neural progenitor like cells were then incubated with neurotrophins in addition to the anti-mitotic agent floxuridine. hNs are then maintained in neurobasal medium and neurotrophins. **c.** Representative images of hNs co-cultured with hpAs and stained for SYNAPSIN1 (red) and MAP2 (green). Cells are counterstained with DAPI (blue). Scale: 100 pixels. Insets: Arrows show SYNAPSIN1 puncta localized on MAP2 positive neurites. **d.** Flowchart of the high content synaptic screening platform. One day after automated seeding of hNs and hpAs (using a liquid handler, day 5 of hN differentiation), a random plate is selected for assessing plating consistency (Columbus script). On day 18, treatments (shown here as small molecules at 3µM) are administered in triplicate for 72 hours using a liquid handler; two plates are used as sentinel or reference. Co-cultures are then stained for synaptic markers using the liquid handler and high-content images are acquired using the Opera Phenix (Perkin Elmer). Data are then processed through CellProfiler and Screener to quantify the number and area of SYNAPSIN1 puncta on MAP2 expressing neurites, the area of MAP2 positive neurites, the number of DAPI positive nuclei and the density of SYNAPSIN1 puncta on MAP2 expressing neurites (Z-score). **e-f.** Acceptance criteria for plating consistency. Plating consistency for each batch is determined using a Columbus script by quantifying detected hN nuclei per well (**e**) as well as the intra-plate covariance (**f**) of the randomly selected plate on day 6. Thresholds for inclusion (red dashed lines) are set as above 4000 detected hN nuclei per well (**e**) and a covariance below 8% across the plate (**f**). **g.** Quantification of the density of SYNAPSIN1 puncta on MAP2 neurites after incubation with SYNAPSIN siRNA versus control conditions. n = 6 wells for each condition; *** p < 0.0001, ANOVA with Dunnett’s post hoc test. Error bars are shown as SEM. See also Figures S1-S5.

Upon hN co-culture with human primary astrocytes (hpAs, ScienCell), we observed a dense synaptic network (**Fig. 1c**). We specifically focused on the presynaptic marker SYNAPSIN1, which is crucial for maintenance, translocation and exocytosis of synaptic vesicle pools, and the cytoskeletal protein MAP2 (microtubule associated protein 2), which is primarily enriched in perikarya and dendrites (Caceres et al., 1984) in addition to DAPI expressing nuclei. We then quantified presynaptic puncta, defined as SYNAPSIN1 puncta assembled on MAP2- expressing neuronal dendrites, which are: (i) robustly captured in our human cellular models; (ii) essential for normal neuronal development with perturbations implicated in disease (Sudhof, 2017; Waites and Garner, 2011); (iii) a prerequisite for the assembly of postsynaptic machinery (Friedman et al., 2000; Rohrbough et al., 2007; Sanes and Lichtman, 2001) and (iv) successfully measured in multiple synaptic screens performed in mouse (Hempel et al., 2011; Spicer et al., 2018). As expected, SYNAPSIN1 appeared in a discrete punctate staining pattern localized on and along the clearly defined MAP2 expressing neurites (**Fig. 1c**). To enhance the resolution of presynaptic puncta and reduce non-specific background for high-content imaging, we included glycine and TrueBlack® Lipofuscin, key reagents for unmasking epitopes and quenching autofluorescence in our immunocytochemistry protocol.

### Establishment of automated pipelines for high-content human presynaptic phenotyping

We next developed a novel and scalable platform for automated quantification of presynaptic puncta in hN and hpA co-cultures (**Fig. 1d**). First, because the number and distribution of synapses in a network is critically dependent upon neuronal density (Cullen et al., 2010), we used an automated liquid handling system (Personal Pipettor, Apricot Designs) to maximize pipetting precision and accuracy and reliably dispense batches of post-mitotic hNs and hpAs across the 60 inner wells of each 96-well plate. To rigorously confirm hN plating consistency, we developed an *in-silico* script using Columbus Acapella software (PerkinElmer; **Supplemental Fig. 2a, b**), quantified DAPI expressing hN nuclei in the co-culture and established key thresholding criteria (*i.e.,* the minimum number of detected hN nuclei per well and the related intra-plate covariance) for each batch of plates (**Fig. 1e, f**). Importantly, hN plating consistency was highly reproducible across independent batches of differentiations (**Fig. 1e, f**).

Second, we developed an automated synaptic quantification pipeline called ALPAQAS (Automated Layered Profiling And Quantitative Analysis of Synaptogenesis) based on image analysis algorithms from the open-source software CellProfiler (Carpenter et al., 2006). After immunostaining, image fields were acquired with a 20x objective with the Opera Phenix high-content screening system (Perkin Elmer), taking 12 fields per well and 5-8 stacks per field at 0.3µm distance (**Supplemental Fig. 3**). We then used ALPAQAS pipelines to quantify MAP2 positive neurites and SYNAPSIN1 puncta parameters based on staining intensity and morphological thresholding (**Supplemental Fig. 3**). The first ALPAQAS pipeline was designed to merge in a single plane (maximum projection) the 5-8 stacks of each channel for each acquired field of a well. The generated output files were the inputs for a second pipeline which corrected for potential illumination variations and translational misalignments, performed morphological reconstruction and de-clustering, and generated a *.csv file reporting at a field level: (1) the area covered by MAP2 positive neurites; (2) the number of SYNAPSIN1 puncta localized on MAP2 positive neurites; (3) the number of DAPI positive nuclei and (4) the area covered by SYNAPSIN1 puncta (**Supplemental Fig. 3**). To validate the specificity of synaptic detection *in vitro*, we assessed presynaptic density (*i.e.,* the number of SYNAPSIN1 puncta localized on the MAP2 positive neurites divided by the area covered by MAP2 positive neurites) using the above pipelines following small interfering RNA (siRNA) perturbation of SYNAPSIN1 (*SYN1*; Accell Dharmacon). Although siRNA-mediated knockdown is heterogeneous, incubating human co-cultures with *SYN1* targeting siRNA for 72 hours significantly reduced presynaptic density compared to *GAPD* and non-targeting siRNA controls, with no or minimal impact on the area covered by MAP2 positive neurites or the total number of DAPI positive neurites in any condition (**Fig 1g; Supplemental Fig. 4a-c**).

Third, to convert the field-level analysis generated by ALPAQAS to well- or condition-level analyses, and to control for potential intra-plate edge and drift effects, we used Genedata Screener software (Genedata). Specifically, the *.csv data file generated from ALPAQAS was imported into Genedata through a high content parser containing three additional quality control steps: (1) data were selected according to their MAP2 field area coverage in comparison to the intra-batch mean MAP2 field area coverage; (2) data from wells of poor quality (fewer than five fields persisting after the first quality control step) were discarded from the analysis and (3) a minimum of two out of three wells per condition (e.g., genotype, treatment) was required by the parser for use in Genedata Screener to aggregate the data from a well-level to a condition-level (**Supplemental Fig. 5**). To control for potential intra-plate edge effects, we included two columns of control wells (0.1% DMSO treated) in each plate. In addition, two plates were assigned as sentinel or reference in each batch of human co-cultures to evaluate and correct eventual anomalies due to intra-batch drift effects. We then leveraged pattern correction algorithms in Genedata Screener to potentially rectify values across plates prior to performing Z-score calculations (Sherman and Bang, 2018; van der Kant *et al*., 2019; Zhang et al., 1999) for synaptic density (**Fig. 1d; Supplemental Fig. 5**).

Collectively, these customized protocols and analysis pipelines establish a novel platform for automated quantification of presynaptic density of hN + hpA co-cultures.

### Primary screening results for 376 small molecules followed by secondary validation reveals six potent small molecules increasing presynaptic density

To validate the robustness of our platform and to uncover modulators of human synaptogenesis, we next assessed the impact of 376 small molecules from a highly selective inhibitor library (SelleckChem, L3500, **Supplemental Table 1**). Here, small molecules were not selected based on known roles in neuronal or synaptic development, but instead from structurally diverse classes of inhibitors targeting kinases, chromatin modifiers and cytoskeletal signaling pathways among others (**Supplemental Table 1, Supplemental Fig. 6a**). Human co-cultures (hNs + hpAs) were treated with small molecules at a concentration of 3µM in triplicate for 72 hours starting on day 18 of hN differentiation (**Fig. 2a**). Importantly, 92.94% of the control and treated wells had a Z-score for synaptic density between -2 and +2, underscoring the reliability and robustness of our automated high-content synaptic screening platform (**Supplemental Fig. 6b**). Thirteen small molecules (3.46% of the library) did not meet the above quality control requirements and seven (1.86% of the library) reduced the density of SYNAPSIN1 puncta on MAP2 neurites by a Z-score ≤ -3 and reduced the area covered by MAP2 positive neurites by a Z-score ≤ -2 (**Fig. 2a; Supplemental Fig. 6c**); as expected these small molecules were known to induce apoptosis or catalytic process of autophagy (Boccadoro et al., 2005; Piva et al., 2008; Reddy et al., 2011). Representative examples of the effects of individual small molecules on presynaptic density are shown in **Fig. 2b**.

**Figure 2:**
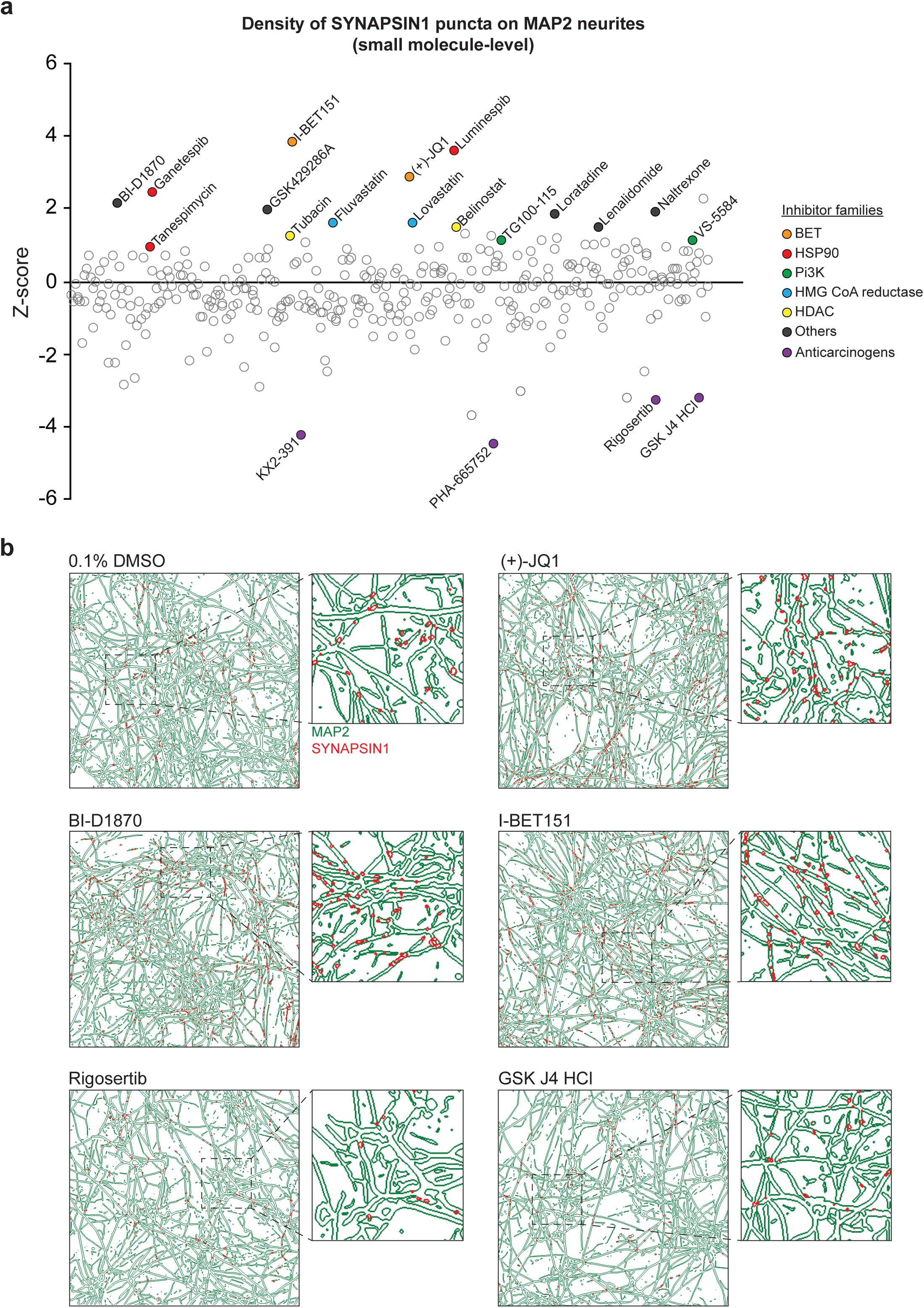
Primary screening results for 376 small molecules. **a.** Z-score values for the density of SYNAPSIN1 on MAP2 expressing neurites for hN + hpA co-cultures incubated with small molecules in triplicate at 3µM for 72 hours starting on day 18 of neuronal differentiation (each circle represents the small molecule level aggregate of the replicates). The effects of each individual small molecule were assessed using our *in silico* pipelines. Small molecules selected for further validation are indicated by colored circles. Those in purple are anticarcinogens. **b.** Representative CellProfiler output images (field-level) of human co-cultures treated with either 0.1% DMSO, (+)-JQ1, BI-D1870, I-BET151, Rigosertib or GSK J4 HCl. Note the increase in SYNAPSIN1 puncta on MAP2-expressing neurites following (+)-JQ1, BI-D1870 and I-BET151 compared to 0.1% DMSO control, and the decrease in SYNPASIN1 puncta following treatment with Rigosertib and GSK J4 HCl. See also Figure S6 and Table S1.

To further validate the top hit classes from our initial small molecule screen, we executed secondary dose-response assays with newly purchased reagents. Specifically, we selected three heat-shock protein 90 inhibitors (Luminespib, Ganestespib and Tanespimycin), three BET inhibitors (I-BET151 and (+)-JQ1 included in the primary screen in addition to Birabresib), two phosphoinositide 3-kinase inhibitors (VS-5584 and TG100-15), two histone deacetylase inhibitors (Tubacin and Belinostat), two HMG CoA reductase inhibitors (Fluvastatin & Lovastatin), a S6 ribosome inhibitor and a ROCK1/ROCK2 inhibitor (BI-D1870 and GSK429286A, respectively), a histamine H1 and an opioid receptor antagonist (Loratadine and Naltrexone, respectively) and a ligand of ubiquitin E3 (Lenalidomide) (**Fig. 2a, b**), for a total of 17 small molecules. We then performed dose-response assays in hN + hpA co-cultures using the automated HP D300e Digital Dispenser (HP, FOL57A) to reliably and randomly dispense at increasing concentrations, nanoliters or microliters of the freshly dissolved small molecules across the 60 inner wells of each 96-well plate and subsequently quantified synaptic density using our previously described ALPAQAS pipelines (**Fig. 3a**).

**Figure 3:**
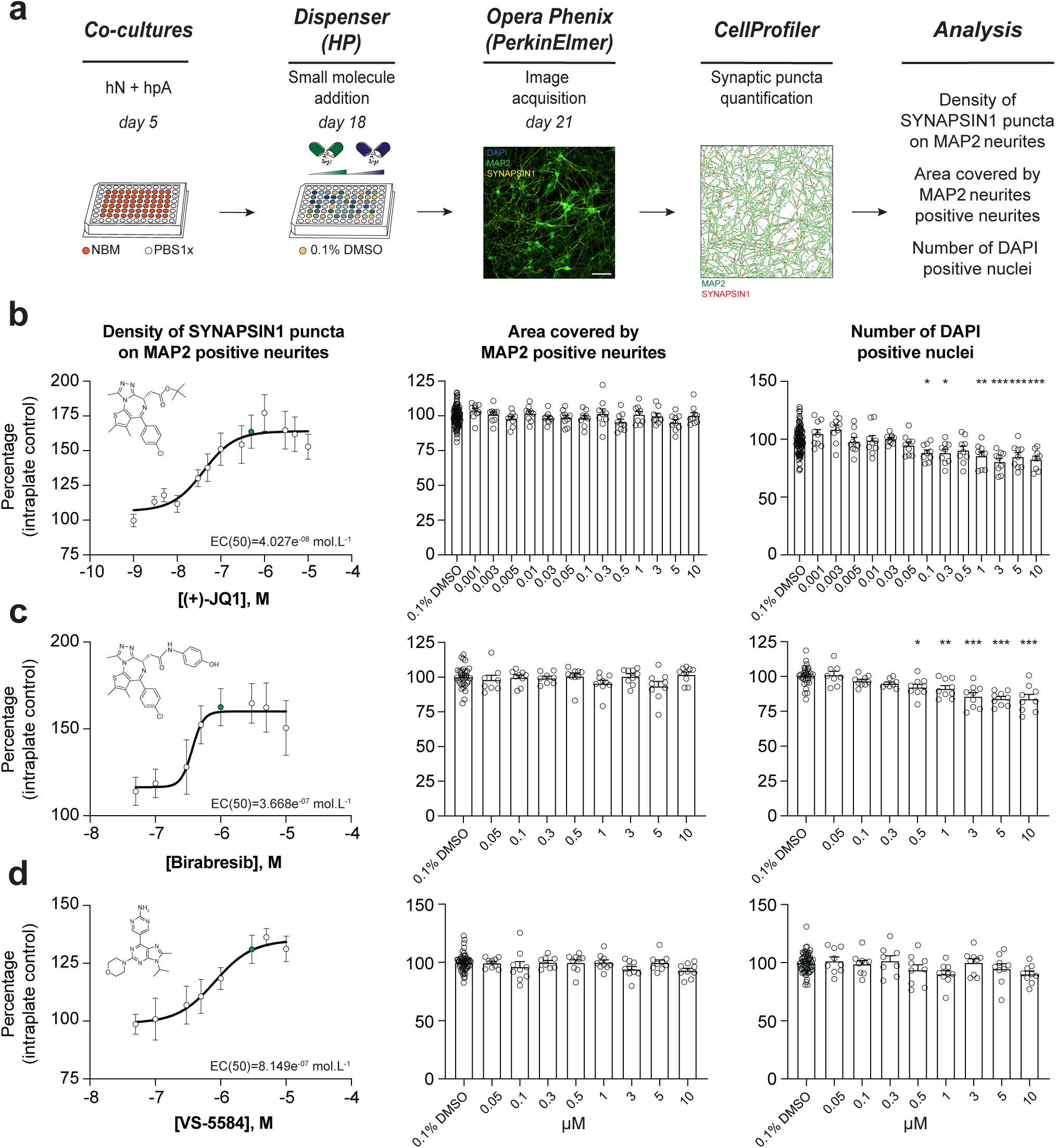
Secondary validation reveals potent small molecules increasing presynaptic density. **a.** Workflow for the dose-response assay. hN + hpA co-cultures were treated on day 18 with selected small molecules at various concentrations or 0.1% DMSO for 72 hours, stained and processed through the synaptic assay. **b**-**d.** Concentration responses for three small molecules (two BET inhibitors ((+)-JQ1, Birabresib) and VS-5584) increasing human presynaptic density (*left*). Green circles indicate the final selected concentration. The impact on the area covered by MAP2 positive neurites (*middle*) and toxicity (number of DAPI positive nuclei; *right*) are also shown. EC(50) and curves were calculated and drawn by PRISM software. Data are quantified by percentage of intraplate control (0.1% DMSO) represented as mean values ± SEM, n = 3 biological replicates, n = 3 technical replicates. *p<0.05, **p<0.01, ***p<0.001; One-way ANOVA with Dunnett’s multiple comparisons test. See also Figures S7-S11.

We found that six out of 17 small molecules tested exhibited a significant dose-dependent increase in presynaptic density compared to DMSO treated controls (**Fig. 3; Supplemental Fig. 7, 8**). Notably, all three BET inhibitors ((+)-JQ1, Birabresib and I-BET151) increased presynaptic density with similar magnitudes of effect (**Fig. 3b-c; Supplemental Fig. 7, 8**). BI-D1870 and GSK429286A also elicited a significant dose-response, indicating that the synaptic connectivity in hNs is modulated through multiple intracellular pathways (**Supplemental Fig. 7, 8**). Indeed, previous studies have shown that ROCK1/ROCK2 inhibition can enhance synapse formation in rodent models (Swanger et al., 2015), consistent with our results from GSK429286A treatment of hN + hpA co-cultures. Finally, VS-5584, an ATP-competitive Pi3K inhibitor with equivalent potency against all human isoforms (α, ≤, ψ and 8) drove a concentration dependent increase in presynaptic density compared to the DMSO treated wells, but not TG100-15, a selective inhibitor of the ψ and 8 isoforms (**Fig. 3d; Supplemental Fig. 7**), consistent with the preferential immune-cell expression of Pi3Kψ and 8 versus the ubiquitous expression patterns of Pi3Kα and ≤ (Winkler et al., 2013). The six validated small molecules showed minimal to no toxicity or effect on neurite outgrowth at lower doses, as assessed by the number of DAPI positive nuclei detected and the area covered by MAP2 positive neurites (**Fig. 3b-d; Supplemental Fig. 8**).

For subsequent experiments, we determined the optimal dosage of each small molecule in hN + hpA co-cultures based upon: (1) a significant increase in synaptic connectivity compared to the DMSO treated wells; (2) a concentration higher than the EC (50); (3) minimal impact on the area covered by MAP2 positive neurites; and (4) minimal or no toxicity (**Fig. 3; Supplemental Fig. 7-10**). We selected 0.5µM for (+)-JQ1, 1µM for Birabresib, 3µM for I-BET151, VS-5584 and BI-D1870, and 5µM for GSK429286A as effective concentrations to increase human presynaptic density. We then derived hNs from an independent hPSC line with the same stable, inducible AAVS1 Ngn2 integration strategy described above, and confirmed that four out of six small molecules, including all three BET inhibitors increased presynaptic density in hN + hpA co-cultures at these specific concentrations (**Supplemental Fig. 11**), further validating our previous findings and indicating that the effects of the selected small molecules were generally reproducible across cell lines. Collectively, these analyses highlight the robustness and the sensitivity of our platform to detect modulators of human synaptogenesis.

### Astrocytes are key regulators of human synapse assembly *in vitro* and response to small molecules

Astrocytes are known to play critical roles in the regulation of neuronal network development (Chung et al., 2015; Perez-Catalan et al., 2021), including the enhancement of presynaptic function (Ullian *et al*., 2001), and we therefore assessed the necessity of hpAs in mediating the effects of the identified small molecules. This was particularly relevant to assess, given that astrocytes are not standardly included in hN screening strategies. Specifically, we tested the impact of the six validated small molecules described above on presynaptic density in the absence of hpAs. hN monocultures were plated, treated, and analyzed using our established pipelines (**Fig. 4a**). While hN monocultures passed all quality control and thresholding criteria (**Supplemental Fig. 12**), none of the six small molecules elicited a significant increase in presynaptic density in this condition (**Fig. 4b**). As expected, the area covered by MAP2 positive neurites and the number of DAPI positive nuclei remained unchanged by small molecule treatment (**Fig. 4c-d**), paralleling the hN + hpA co-culture condition (**Fig. 3**). Importantly, experiments assessing the effect of the six validated small molecules on an independent hPSC line using hN + hpA co-cultures (**Supplemental Fig. 11**) were used as a positive control and performed side-by-side experiments assessing the effect of the six validated small molecules on hN monocultures (**Fig. 4**). Small molecule treatment also did not significantly affect the level of SYNAPSIN1 protein expression in either the hN monoculture or hN + hpA co-culture conditions as assessed by western blot analysis (**Fig. 4e-f**), indicating that the differences observed between hN monocultures and hN + hpA co-cultures after small molecule addition were more likely due to changes in SYNAPSIN1 protein localization as opposed to overall protein abundance.

**Figure 4:**
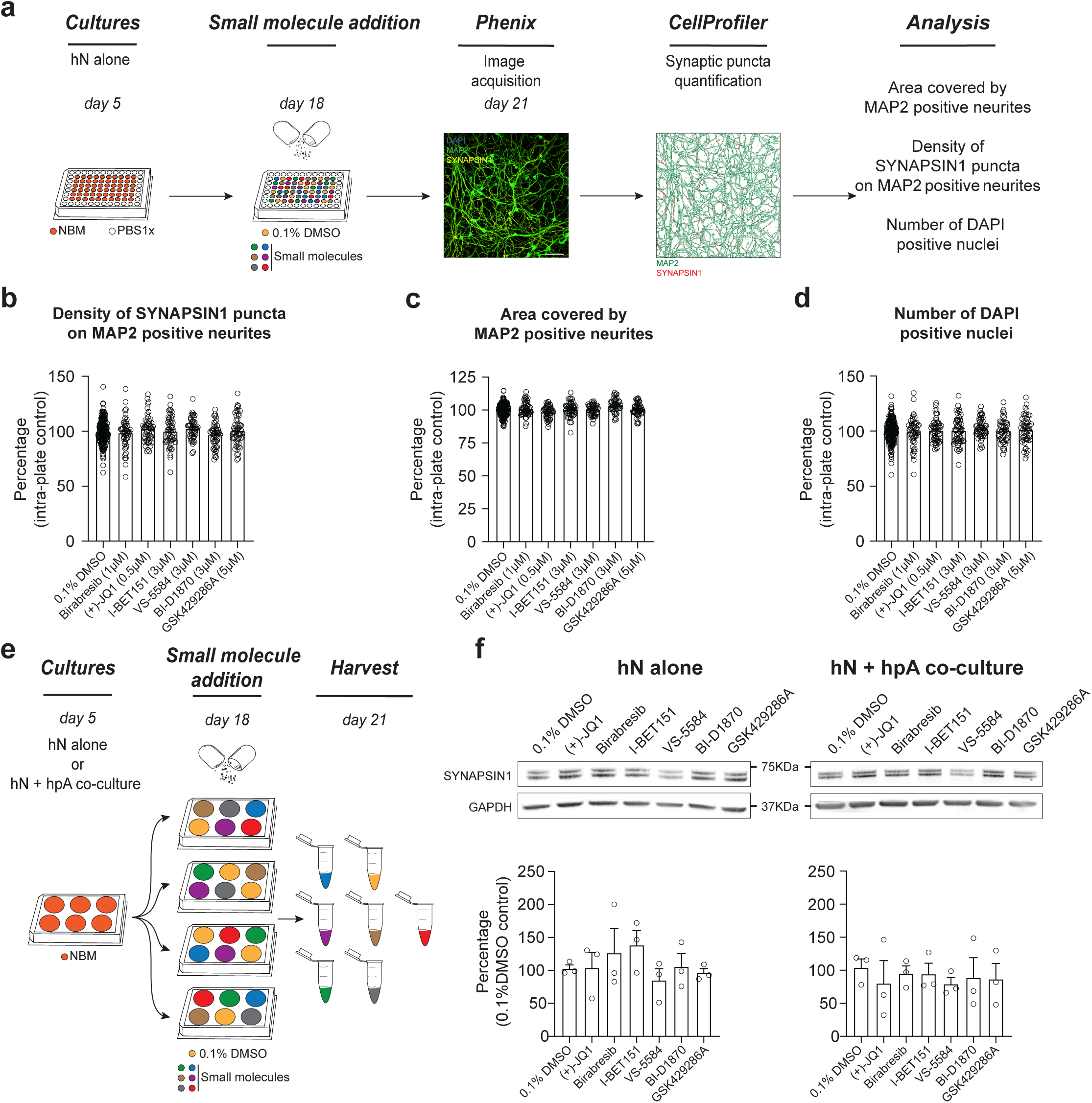
Astrocytes play a critical role in mediating response to small molecules. **a.** Workflow for small molecule validation in the absence of hpAs. **b**-**d.** In the absence of hpAs, none of the selected small molecules affected SYNAPSIN1 density of hNs, in contrast to the co-culture condition. The area covered by MAP2 positive neurites and cell viability were also not impacted as compared to intra-plate DMSO controls. **e.** Schematic for the preparation of the immunoblot samples. **f.** Representative images of immunoblots and quantification of SYNAPSIN1 normalized to GAPDH and presented as a percentage of DMSO control in both hN alone and hN + hpA co-culture conditions. Note SYNAPSIN1 protein expression levels were not impacted in any condition. n = 3 biological replicates, n ;: 3 technical replicates (**f)**. See also Figure S12.

The addition of hpAs also had a significant effect on synapse development in the absence of small molecules, including a significant increase in SYNAPSIN1 presynaptic density as well as in the size of individual SYNAPSIN1 puncta with a modest reduction in the area covered by MAP2 positive neurites (**Fig. 5a-e**). Indeed, the addition of hpAs roughly doubled both the density and size of SYNAPSIN1 puncta on MAP2 positive neurites compared with hNs alone (**Fig. 5c-d**). Collectively, our results indicate that hNs can form quantifiable presynaptic puncta which can be reliably captured by our platform in the absence of astrocytes. However, hpAs contributed to the assembly or localization of presynaptic machinery, underscoring the relevance of astrocytes in human synaptic screening applications. hpAs were therefore included in all subsequent experiments.

**Figure 5:**
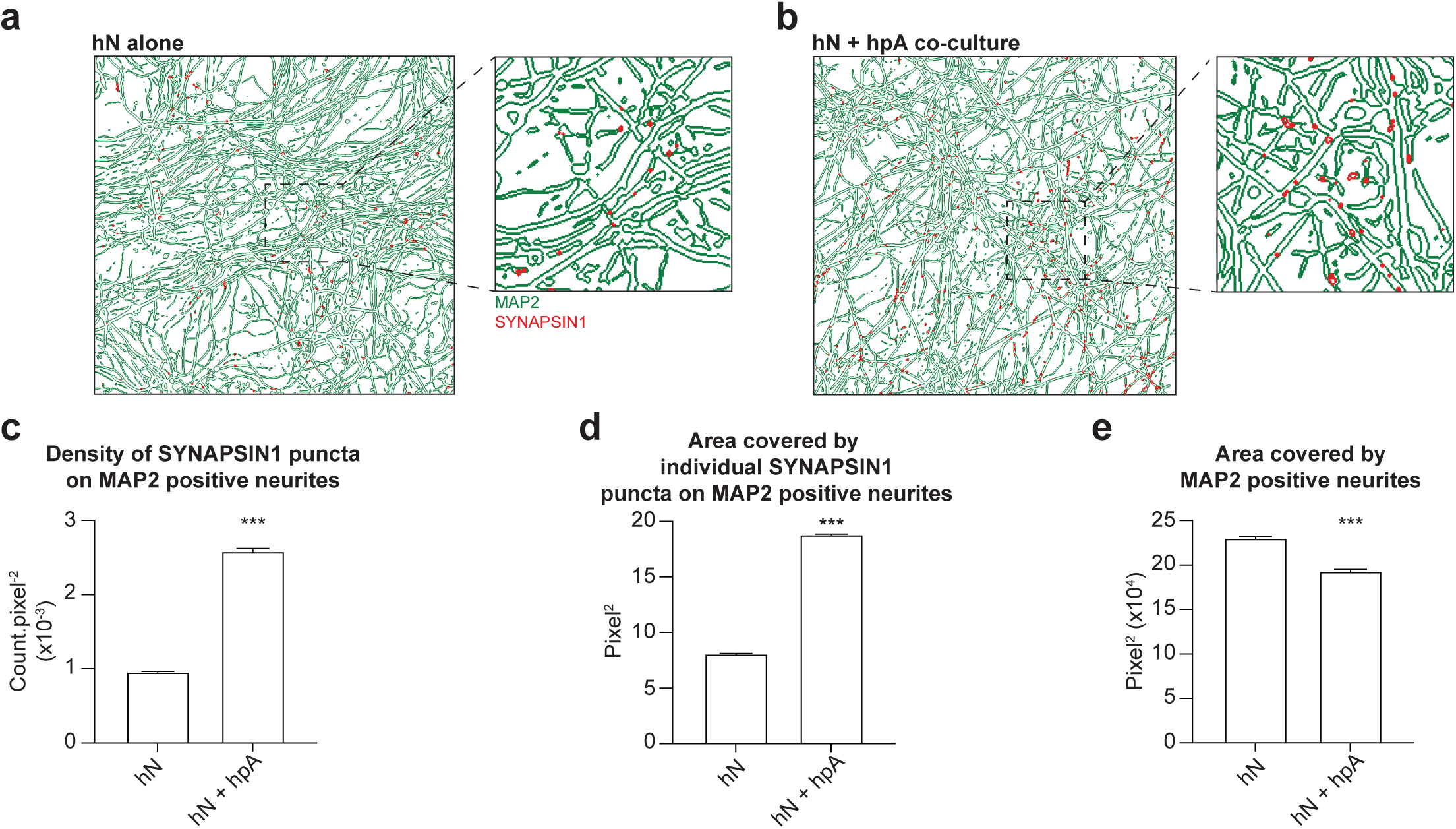
Astrocytes are key regulators of human synapse assembly *in vitro*. **a-b.** Representative CellProfiler output images (field-level) of hN monocultures and hN + hpA co-cultures. **c-e.** Measurements comparing the density of SYNAPSIN1 puncta on MAP2 positive neurites (**c**), the area covered by individual SYNAPSIN1 puncta (**d**) and the area covered by MAP2 positive neurites (**e**) between hN monocultures and hN + hpA co-cultures. Data are represented as mean values ± SEM, n = 3 biological replicates, n = 8 technical replicates (**c-e)**; n > 1,000 fields and n > 50,000 SYNAPSIN1 puncta for each condition (**c**-**e**). ***p<0.001; Kolmogorov-Smirnov unpaired t-test.

### Multiple BET inhibitors enhance synaptic gene expression

Given that BET inhibitors were the most prominent hit class identified in our small molecule screen, with three independent BET inhibitors increasing presynaptic density in two independent hPSC lines, we sought to confirm a role for BET proteins in human synaptic development and further validate the results of our synaptic assay. The BET family of chromatin readers, which includes BRD2, BRD3, BRD4 and BRDT, bind acetylated lysine residues on histone proteins as well as transcription factors to mediate gene expression. Through unbiased screening, (+)-JQ1 was identified as a positive modulator of human neurogenesis (Li et al., 2016). However, in adult mice, BET inhibition through (+)-JQ1 was shown to reduce synaptic gene expression and impair memory consolidation (Korb et al., 2015), while BET inhibition through I-BET858 in mouse primary neurons reportedly led to decreased expression of neuronal differentiation and synaptic genes (Sullivan et al., 2015). Importantly, region-specific differences in response to (+)-JQ1 have been identified in the rodent brain, particularly in the context of dendritic spine density, suggesting brain cell type may be a major determining factor in the expression of these phenotypes (Wang et al., 2021b). Moreover, a recent study found sex divergent effects of Brd4 on cellular and transcriptional phenotypes in both human and mouse (Kfoury et al., 2021), further supporting the hypothesis that the impacts of BET inhibition are highly context dependent.

We therefore performed global transcriptional analyses on both (+)-JQ1 and Birabresib treated hN + hpA co-cultures compared with DMSO treated controls (**Fig. 6a**) to independently assess their roles in synaptic development. (+)-JQ1, Birabresib or DMSO was added on day 18 *in vitro* for 72hrs, paralleling the approach used in our small molecule screen. With an adjusted p-value cut-off of 0.05 and log_2_ fold change cut-offs of ≤-1 and ≥ 1, we identified a comparable number of significantly differentially expressed genes (DEGs) after (+)-JQ1 and Birabresib treatment compared to DMSO control (**Fig. 6b-d; Supplemental Tables 2, 3**). A majority of DEGs were downregulated after BET inhibition (**Fig. 6b-d**), consistent with the known roles of BET proteins in transcriptional activation, and were also shared between the (+)-JQ1 and Birabresib treatment conditions (n=2,368; **Fig. 6e**). Given that both BET inhibitors increased presynaptic density (**Fig. 3b,c**), we focused on the set of DEGs that were shared between (+)- JQ1 and Birabresib treatment conditions for the remainder of our transcriptional analyses. Using Ingenuity Pathway analysis (IPA), we confirmed that BRD4 (p=1.61×10^-17^), (+)-JQ1 (p=0.000206) and Birabresib (p=4.59×10^-18^) were all strongly predicted to be upstream regulators of the shared DEGs (**Supplemental Fig. 13**). Signaling pathways such as “DNA methylation and transcriptional repression” and “axonal guidance” were enriched in the upregulated DEGs, while signaling pathways such as “wound healing” and “neuroinflammation” enriched in the downregulated DEGs (**Fig. 6f**).

**Figure 6:**
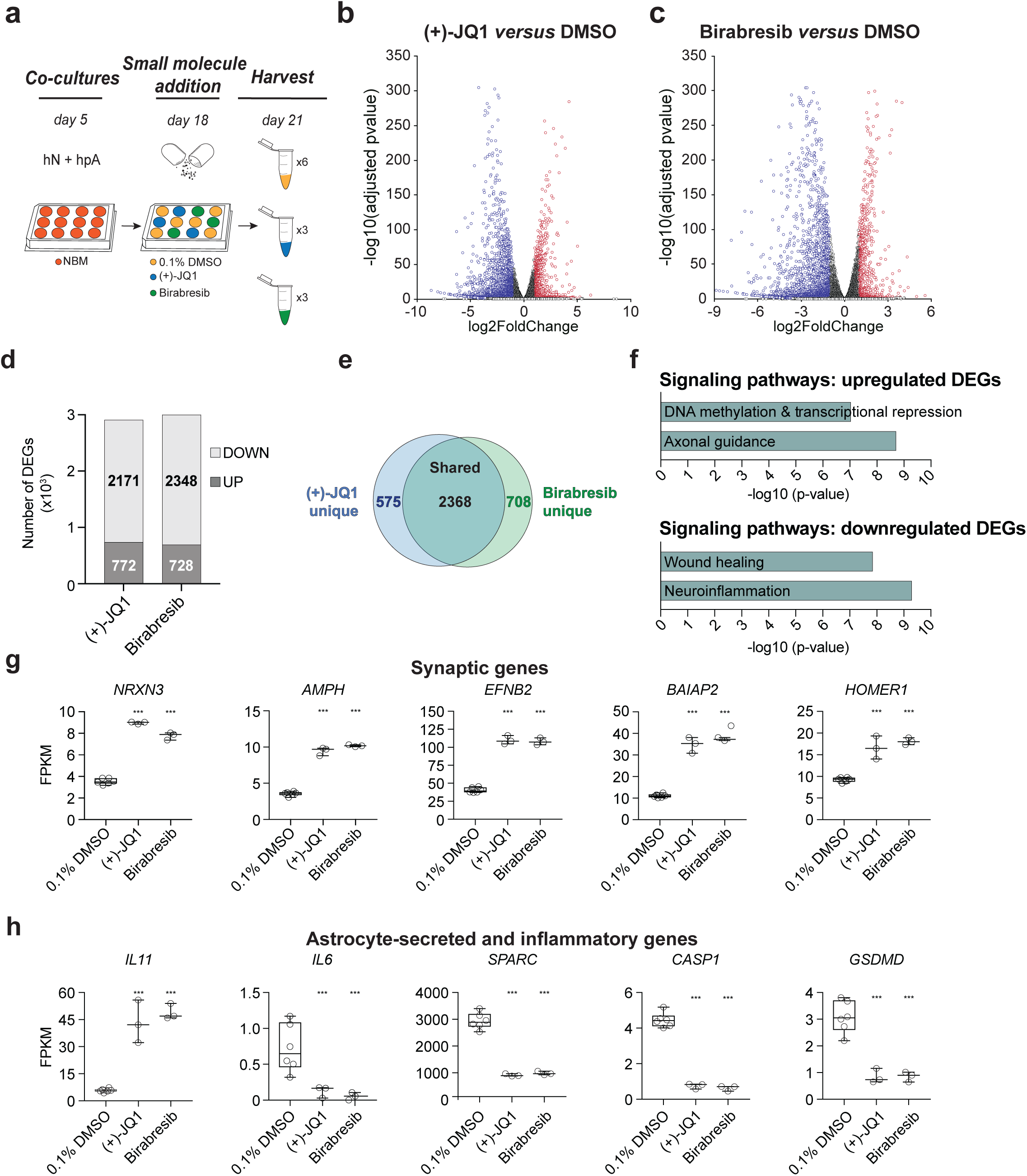
Multiple BET inhibitors enhance synaptic gene expression programs. **a.** Schematic of mRNA-seq experiment using three to six replicates per treatment condition. hN + hpA co-cultures were treated with 0.1% DMSO, 0.5µM (+)-JQ1 or 1µM Birabresib for 72hrs prior to harvesting. **b-c.** Volcano plots for (+)-JQ1 versus DMSO control (**b**) and Birabresib versus DMSO control (**c**). Log_2_ fold change is shown on the x-axis, with the -log_10_ of the adjusted p-value shown on the y-axis. Positive fold change reflects an increase in the treatment condition relative to DMSO control. **d.** Bar graph showing numbers of significantly differentially expressed genes (DEGs) between treatment and control conditions using an adjusted p-value cutoff of 0.05 and log_2_ fold change cutoff of +/-1. Significantly downregulated genes are shown in light gray and upregulated genes are shown in dark gray. **e.** Overlap (shown as number of genes) between DEGs identified with (+)-JQ1 treatment (blue) and DEGs identified with Birabresib treatment (green). **f.** Select Canonical Pathways identified by IPA for DEGs upregulated in both (+)-JQ1 and Birabresib treatment conditions (top) or downregulated in both (+)-JQ1 and Birabresib treatment conditions (bottom). The x-axis shows -log_10_ of the p-value for each pathway analysis term. **g-h.** Graphs of transcript expression values for pre- and post-synaptic genes (**g**) and astrocyte-secreted and inflammatory genes (**h**). Significance was calculated by Benjamini-Hochberg adjusted Wald test as part of the DEseq2 RNA-seq experiment. For all figure panels, significance is indicated by *** p≤0.0005 relative to controls. See also Figure S13 and Tables S2-S3.

Closer examination of individual shared DEGs revealed significant upregulation at the transcriptional level of both pre- and post-synaptic genes following BET inhibition, including the presynaptic cell adhesion molecule *Neurexin 3 (NRXN3),* the synaptic vesicle gene *Amphiphysin (AMPH),* the transmembrane Eph receptor ligand *Ephrin-B2 (EFNB2)* involved in AMPAR stabilization, and the postsynaptic scaffolding factors *BAR/IMD Domain Containing Adaptor Protein 2 (BAIAP2*) and *Homer Scaffold Protein 1* (*HOMER1)* (**Fig. 6g**). Importantly, changes in gene transcription were highly concordant between both (+)-JQ1 and Birabresib treatment conditions, with similar magnitudes of effect (**Fig. 6g**). Indeed, (+)-JQ1 and Birabresib treatment each roughly doubled the expression of these pre- and post-synaptic genes.

Consistent with the pathway analyses, we also observed modulation of the expression of several astrocyte-secreted factors including upregulation of the anti-inflammatory factor *Interleukin 11 (IL11)*, downregulation of the pro-inflammatory factor Interleukin 6 (*IL6)* and downregulation of the synapse disassembly factor *Secreted Protein Acidic And Cysteine Rich (SPARC)* (**Fig. 6h**). Combined with downregulation of Caspase 1 (*CASP1) and Gasdermin D (GSDMD)* (**Fig. 6h**), these results point to a general anti-inflammatory response following BET inhibition. Indeed, *CASP1, GSDMD* and *IL6* have all previously been shown to be downregulated by (+)-JQ1 treatment, consistent with the known role of BET proteins in promotion of the inflammatory response (Wang *et al*., 2021b; Zhou et al., 2019). Importantly, changes in gene transcription were again highly concordant between both (+)-JQ1 and Birabresib treatment conditions, with similar magnitudes of effect (**Fig. 6h**).

Collectively, these data indicate that BET inhibition leads to upregulation of synaptic gene expression programs, consistent with our screening results showing an increase in presynaptic density and supporting a role for BET proteins in synapse assembly. These results support the robustness of our platform for detecting human synaptic modulators *in vitro*.

## DISCUSSION

Despite overwhelming evidence implicating synaptic dysfunction in complex human disease, and clear species-specific differences in certain aspects of synaptic biology, human neurons and astrocytes have yet to be widely employed in studies of synaptic mechanisms. We therefore established a novel, automated, high-content synaptic platform using human neurons and astrocytes to facilitate dissection of human synaptic mechanisms and drug discovery efforts. Specifically, we: (i) generated stable iNgn2-hPSC lines through TALEN editing to produce large, reproducible batches of post-mitotic glutamatergic hNs; (ii) optimized hN + hpA co-culture and immunocytochemistry conditions for reliable quantification of synaptic markers on an automated imaging platform; (iii) employed liquid handling systems to standardize the dispensing steps and maximize accuracy and (iv) developed *in silico* automated pipelines for rigorous assessment of plating consistency and quantification of human synaptic connectivity. Through unbiased screening of 376 small molecules, we generated proof-of-concept data that our platform works robustly to identify synaptic modulators. Our results were also generally reproducible across independent hPSC lines. Given the known molecular and functional heterogeneity across individual hPSC lines (Kajiwara et al., 2012; Kilpinen et al., 2017; Schwartzentruber et al., 2018), we would not expect identical results or magnitudes of effect, but our analyses support the robustness of our assay to detect differences in independent hPSC lines.

Importantly, the results from hNs co-cultured with hpAs did not correlate with the results from hNs cultured alone, underscoring the importance of including astrocytes in human synaptic assays. Indeed, our positive hits required the presence of hpAs to achieve a significant change in presynaptic density, supporting the relevance of non-cell autonomous factors in synapse assembly. In line with this observation, upon hpA co-culture we observed a two-fold increase in presynaptic density and the area of individual SYNAPSIN1 puncta compared with hN monocultures. While previous studies have shown that inclusion of rodent astrocytes with hNs can significantly increase physiological and/or morphological maturation (Johnson et al., 2007; Nehme *et al*., 2018; Tang et al., 2013), the underlying mechanisms remain incompletely resolved. Studies examining the effects of physical astrocyte contact versus astrocyte conditioned media on hNs report that physical contact has a much more potent impact on synapse development compared with conditioned media (Johnson *et al*., 2007; Tang *et al*., 2013), suggesting that direct physical contact between hNs and hpAs may be particularly important. Our results imply that regulators of synapse formation may be missed in assays performed in the absence of astrocytes. Given that the inclusion of astrocytes more accurately mimics the *in vivo* synaptic milieu, and that astrocytes are not standardly included in hN screening strategies, it will be critical to consider this cell type in future human synaptic screening experiments.

Given that three separate BET inhibitors scored in our assay in two independent hPSC lines, our results also support a role for BET proteins in human synaptic development. While we were unable to separate hN and hpA transcriptomes from the co-culture (and the co-culture was required to see the effects of BET inhibition), we observed modulation of synaptic machinery in our global transcriptional analyses, consistent with our screening results. These data support a model whereby BET inhibition either leads to increased expression of synapse associated genes, or to the enhancement of presynaptic puncta which then triggers the expression of additional synaptic components. Indeed, BET protein expression is downregulated throughout neuronal differentiation, with knockdown of BET proteins sufficient to enhance differentiation (Li *et al*., 2016) consistent with a role for BET proteins in negatively regulating neuronal development. These results contrast with a subset of data generated in adult mouse brain and are thus likely to be dependent upon factors such as brain cell type, developmental stage, and timing of drug treatment. Resolving potential neurological benefits and/or consequences of BET inhibition in different contexts including in diverse brain cell types and developmental stages is essential, as these drugs are currently in use in clinical trials for cancer and are proposed for the treatment of developmental as well as degenerative diseases (Korb *et al*., 2015; Wang et al., 2021a).

While BET proteins have well-described roles in the inflammatory response, little is known about their role in astrocytes specifically. Future studies will be required to determine how BET inhibition impacts astrocytes, and how this may in turn impact neuron function. Our results are consistent with BET inhibition enhancing synapse assembly and impacting both neuron and astrocyte transcriptomes, but it remains to be determined whether BET inhibition of astrocytes has a direct impact on synapse formation, or whether BET inhibition primarily acts on neurons to impact synapse assembly. Overall, we speculate that hpAs provide critical synaptic support for hNs in our assay which then facilitates the health or receptiveness of hNs for further synaptic enhancement, given the disparities in SYNAPSIN1 parameters between hN monocultures and hN + hpA co-cultures, as well as the lack of effect of small molecules on hN monocultures.

Our robust, scalable and accessible platform provides several key advances for the field. Specifically: (i) we generated the first automated and quantitative high-content synaptic phenotyping platform for human neurons and astrocytes, which can now be leveraged for study of basic human synaptic mechanisms, dissection of complex human disease and drug discovery efforts; (ii) our results demonstrate that the stable Ngn2 integration strategy for scaled generation of hNs from hPSCs is well-suited for screening applications; (iii) our platform is compatible with both small-molecule as well as siRNA based manipulations; (iv) our platform captures key parameters of human synaptic connectivity including the number and the area of individual presynaptic contacts as well as the coverage of MAP2 positive neurites and overall cell viability; (iv) while compatible with analyses of hN monocultures, our analyses underscore the relevance of astrocytes when studying human synaptic mechanisms and drug response and (v) different human genetic backgrounds can be employed in our platform to dissect disease mechanisms. Future studies will focus on the incorporation of additional brain cell types to increase the complexity in our 2D co-culture systems, probing additional small molecule classes to explore novel human synaptogenic factors and improving the fidelity of astrocyte differentiation protocols *in vitro* to facilitate cell-type specific genetic manipulations.

There are also several limitations to our approach: (i) our platform was specifically designed for use with human neurons and astrocytes and alterations to the neuronal differentiation paradigms or species, or the addition of other brain cell types may require modest customization steps; (ii) our platform is based on high-content imaging, and neurophysiological function must be assessed using separate methodology, likely at lower throughput; (iii) while the time-points of manipulations and analyses can be altered in our assay, longitudinal studies in the same cells are not possible unless suitable genetically encoded fluorescent markers are employed, and (iv) our platform is designed for 2D culture to allow for precise control over cell densities and ratios but does not fully reflect the structural complexity present in 3D systems (e.g., spheroids, organoids).

## MATERIALS AND METHODS

### Stem cell culture

The XY human ESC line WA01 (H1) and the XY human iPSC line DS2U were commercially obtained from WiCell Research Institute (Thomson et al., 1998; Weick et al., 2013) (www.wicell.org). Stem cell culture was carried out as previously described (Hazelbaker et al., 2020; Hazelbaker et al., 2017). In brief, stem cells were grown and maintained in StemFlex medium (Gibco, A3349401) on geltrex pre-coated plates (Life Technologies, A1413301) under standard conditions (37°C, 5% CO_2_). Cells were passaged using TrypLE Express (Life Technologies, 12604021). All cell lines underwent QC testing to confirm normal karyotypes, absence of mycoplasma, expression of pluripotency markers and tri-lineage potential. G-band karyotyping analysis was performed by Cell Line Genetics.

### Generation of inducible Ngn2 system

TALENs (AAVS1-TALEN-L and AAVS1-TALEN-R; Addgene, 59025/59026)(Gonzalez et al., 2014) were used to target the first intron of the constitutively expressed gene *PPP1R12C* at the AAVS1 locus with the pAAVS1-iNgn2-Zeo plasmid containing TetO-Ngn2-P2A-Zeo and CAG-rtTA. In brief, hPSCs were dissociated into single cell suspension with TrypLE (Gibco, 12604-021). 2.5×10^6^ cells were resuspended in 120µL of R Buffer (ThermoFisher Scientific, MPK10096) and mixed with 1.5 µg each of AAVS1*-*TALEN-L and AAVS1-TALEN-R, and 10 µg of pAAVS1-iNgn2-Zeo plasmid. Cells were electroporated using the Neon Transfection electroporation system (ThermoFisher Scientific, MPK10096) at 1050 V, 30 ms and 2 pulses, and plated on a 10-cm plate. 24h after electroporation and indefinitely, the cells were selected with geneticin (50µg.mL^-1^, Life Technologies, 10131035). At the fifth day of selection, 2×10^4^ cells were plated on a 10-cm plate for clonal selection. After colony formation, colonies were picked and transferred to a 96-well plate for genomic DNA extraction and PCR analysis of plasmid integration.

### Genomic DNA isolation and genotyping PCR

Genomic DNA (gDNA) from the iNgn2-H1 cell line was extracted from hPSCs with the DNeasy Blood and Tissue kit according to manufacturer’s instructions (Qiagen, 69504). PCR of the gDNA was performed with the primer pair forward 5’-AGGAAATGGGGGTGTGTCAC-3’ (in the AAVS1 locus) and reverse 5’- GAGCTCCTCTGGCGATTCTC-3’ (in the Ngn2 DNA sequence).

### Human neuron generation

Human neurons were generated as previously described (Nehme *et al*., 2018; Zhang *et al*., 2013). In brief, on day 0, hPSCs were differentiated in N2 medium [500 mL DMEM/F12 (1:1) (Gibco, 11320-033)], 5 mL Glutamax (Gibco, 35050-061), 7.5 mL Sucrose (20%, SIGMA, S0389), 5 mL N2 supplement B (StemCell Technologies, 07156)] supplemented with SB431542 (10 µM, Tocris, 1614), XAV939 (2 µM, Stemgent, 04-00046) and LDN-193189 (100 nM, Stemgent, 04-0074) along with doxycycline hyclate (2 µg.mL^-1^, Sigma, D9891) and Y27632 (5 mM, Stemgent 04-0012). Day 1 was a step-down of small molecules, where N2 medium was supplemented with SB431542 (5 µM, Tocris, 1614), XAV939 (1 µM, Stemgent, 04-00046) and LDN-193189 (50 nM, Stemgent, 04-0074) with doxycycline hyclate (2 µg.mL^-1^, Sigma, D9891) and Zeocin (1 µg.mL^-1^, Invitrogen, 46-059). On day 2, N2 medium was supplemented with doxycycline hyclate (2 µg.mL^-1^, Sigma, D9891) and Zeocin (1 µg.mL^-1^, Invitrogen, 46-059). Starting on day 3, cells were maintained in Neurobasal media [500 mL Neurobasal (Gibco, 21103-049), 5 mL Glutamax (Gibco, 35050-061), 7.5 mL Sucrose (20%, SIGMA, S0389), 2.5 mL NEAA (Corning, 25-0250Cl)] supplemented with B27 (50x, Gibco, 17504-044), BDNF, CTNF, GDNF (10 ng.mL^-1^, R&D Systems 248-BD/CF, 257-NT/CF and 212-GD/CF) and doxycycline hyclate (2 µg.mL^-1^, Sigma, D9891). From day 4 to day 5, Neurobasal media was complemented with the antiproliferative agent floxuridine (10 µg.mL^-1^, Sigma Aldrich, F0503-100MG).

### Human primary astrocytes

Human primary cortical astrocytes (hpA) were obtained from ScienCell Research Laboratories (1800) and cultured according to the manufacturer’s instructions.

### mRNA sequencing and analysis

Three to six biological replicates of hN + hpA co-cultures per condition (72hr DMSO treated, 72hr (+)-JQ1 treated, 72hr Birabresib treated) were harvested in RLTplus Lysis buffer (Qiagen, 1053393). Total RNA was isolated using the RNeasy micro/mini plus kit (Qiagen, 74034). Libraries were prepared using Roche Kapa mRNA HyperPrep strand specific sample preparation kits from 200ng of purified total RNA according to the manufacturer’s protocol using a Beckman Coulter Biomek i7. The finished dsDNA libraries were quantified by Qubit fluorometer and Agilent TapeStation 4200. Uniquely dual indexed libraries were pooled in equimolar ratio and subjected to shallow sequencing on an Illumina MiSeq to evaluate library quality and pooling balance. The final pool was sequenced on an Illumina NovaSeq 6000 targeting 30 million 100bp read pairs per library. Sequenced reads were aligned to the UCSC hg19 reference genome assembly and gene counts were quantified using STAR (v2.7.3a)(Dobin et al., 2013). Differential gene expression testing was performed by DESeq2 (v1.22.1)(Love et al., 2014). RNAseq analysis was performed using the VIPER snakemake pipeline (Cornwell et al., 2018). Library preparation, Illumina sequencing and VIPER workflow were performed by the Dana-Farber Cancer Institute Molecular Biology Core Facilities.

### Automated cell plating

hNs and hpAs were harvested with Accutase (Innovative Cell Technology, Inc., AT104-500), quenched in Neurobasal media, spun 5 minutes at 1000rpm at room temperature, passed through a 40µm filter and counted using the Countess Automated Cell Counter (ThermoFisher Scientific, AMQAX1000). Cells were mixed in NBM media to reach a seeding density of 40,000 neurons.cm^-2^ and 100,000 astrocytes.cm^-2^ per well. A liquid handling dispenser (Personal Pipettor, ApricotDesigns) was used to uniformly plate the cell mixture in the geltrex-coated 60-inner wells of 96-well plates (PerkinElmer, CellCarrier-96, 6005558).

### Small molecule dilution and addition

The Selleck library (L3500) consisted of 8 x 96-well plates containing 376 small molecules at 10mM in DMSO (100%). On the day of small molecule addition, the relevant plate was diluted with an automatic liquid handling dispenser (ApricotDesigns, Personal Pipettor). Each diluted plate was screened on the same day, with each small molecule tested against the cell type of interest in three different 96-well plates (in triplicate) at 3 µM.

### siRNA-mediated knockdown

Human SYNAPSIN1 and control siRNA (Accell Dharmacon, A-12362-16-0005 and K-005000-R1-01) were aliquoted (100µM) and stored at -20°C according to manufacturer’s recommendations. siRNA (1µM) in Accell Delivery Media (Accell Dharmacon, B-002222-UB-100) was added to human co-cultures on day 18 of hN differentiation for 72hrs followed by fixation and analysis.

### Immunocytochemistry

Immunofluorescence was performed using an automatic liquid handling dispenser (ApricotDesigns, Personal Pipettor). Cells were washed abundantly in 1x PBS, fixed for 20 minutes in PFA (4%, Electron Microscopy Sciences, 15714-S) plus Sucrose (4%, SIGMA, S0389), washed abundantly in 1x PBS, permeabilized and blocked for 20 minutes in Horse serum (4%, ThermoFisher, 16050114), Triton X-100 (0.3%, SIGMA, T9284) and Glycine (0.1M, SIGMA, G7126) in 1x PBS. Primary antibodies were then applied at 4°C overnight in 1x PBS supplemented with Horse serum (4%, ThermoFisher, 16050114). The following synaptic antibodies were used: Rabbit anti-human SYNAPSIN1 (1:1000, Millipore, AB1543), Chicken anti-human MAP2 (1:1000, Abcam, ab5392). After abundant washes with Triton X-100 (0.3%, SIGMA, T9284) in 1x PBS, cells were exposed for 1h at RT to secondary antibodies: Goat anti-chicken AlexaFluor 488 (1:1000, ThermoFisher, A21131), Donkey anti-rabbit AlexaFluor 555 (1:1000, ThermoFisher, A31572), as well as DAPI (1:5000, ThermoFisher Scientific, D1306) and TrueBlack (1:5000, Biotium, 23007) in 1x PBS supplemented with Horse serum (4%, ThermoFisher, 16050114). Finally, cells were abundantly washed with 1x PBS and stored at 4°C.

### Immunoblotting

Cell lysates were made from iNgn2-H1 hNs monocultures or from hN + hpA co-cultures (40,000 neurons.cm^-2^ and 100,000 astrocytes.cm^-2^ per well) on 6-well plates. Medium was refreshed once per week. Cells were treated with the selected small molecules at the optimal concentration for 72 hours starting on day18 of hN differentiation. Cells were abundantly washed with PBS1x before lysis in RIPA buffer (Life Technologies, 89901) supplemented with protease and phosphatase inhibitor (Thermo Scientific, 88669). Before blotting, samples were triturated with repetitive up and down using U-100 Insulin Syringe (B-D, 329461), centrifuged at 10,000g for 15 minutes at 4°C and the supernatants were collected. Proteins were denatured by boiling at 95°C for 10 mins and subjected to a Pierce^TM^ BCA protein assay (Life Technologies, 23227) for determination of the protein concentration. For each sample the same amount of total protein extracted (5-10µg) was loaded and separated by SDS-PAGE. The proteins were then transferred to a PVDF membrane (Bio-Rad, 1704156), blocked in 5% Difco^TM^ Skim Milk (BD, 232100) in TBS-Tween 20 (1/1000; Sigma-Aldrich, T5912-1L and P9416-50ML) before blotting with the primary antibodies: Rabbit anti-human SYNAPSIN1 (1:1000, Millipore, AB1543) and Mouse anti-human GAPDH (1:1000, Millipore, MAB374). For visualization Donkey anti-Mouse IRDye® 800CW (1:5000, LI-COR, 926-32212) and Donkey anti-Rabbit IRDye® 680RD (1:5000, LI-COR, 926-68073) were used before detection with Odyssey® DLx imaging system (LI-COR, 9140).

### Image acquisition and analysis paradigms

#### Plating consistency

24 hours after co-culture, a random 96-well-plate was fixed and stained. 2 channels (DAPI, MAP2), 4 stacks.field^-1^, each stack separated by 0.5µm, 25 fields.well^-1^, 60 wells.96-well plate^-1^ were acquired at 20x (Plan Apo λA, NA 0.75, air objective, Nikon) with a high content screening confocal microscope (ImageXpress Micro 4, Molecular Devices). Acquired images were transferred and stored on a Columbus (PerkinElmer) server. A Columbus Acapella software (PerkinElmer) algorithm was designed to discriminate hN nuclei from hpA nuclei. DAPI and MAP2 channels of each field were merged in a single plane to create maximum projection images. DAPI projected images were filtered using a Gaussian method and the nuclei population was detected based on the C method (diameter > 20µm). Intensity and morphological properties of each identified nuclei were calculated. Among all nuclei, the hN nuclei were identified according to their intensity, contrast as well as area, roundness and distinct MAP2 positive soma (**Supplemental Fig. 2b**). This script allowed quantification of the mean number (and the standard deviation) of hNs identified in each well, and the coefficient of variance (%) per plate. Thresholding for plating consistency was set above 4,000 hNs per well and a covariance below 8% per plate.

#### Synapse detection

Three channels (DAPI, MAP2, SYNAPSIN1), 5-8 stacks.field^-1^ at 0.3µm distance, 12 fields.well^-1^, 60 wells.96-well plate^-1^ were acquired with a 20x objective (NA 1.0, water objective) with a high-content screening confocal microscope (Opera Phenix, PerkinElmer). Image analysis pipelines were built using the open-source CellProfiler 3.1.5 software (www.cellprofiler.org)(Carpenter *et al*., 2006) Each of the twelve fields per well were analyzed independently. The first pipeline was designed to merge in a single plane (maximum projection) the 5-8 stacks of the raw image for each channel. The output files generated by the first pipeline were the input files for a second pipeline. The second pipeline was developed to report different features in MAP2 and SYNAPSIN1 channels. First, translational misalignment was corrected by maximizing the mutual information of the DAPI and MAP2 channels. The same alignment measurements obtained from the first two input images were applied for the SYNAPSIN1 channel. Illumination correction functions were created by averaging each pixel intensity of all images from each channel across each plate then smoothing the resulting image with a Gaussian filter and a large filter size of 100 x 100 pixels. A rescale intensity function was used to stretch each image of each channel to the full intensity range (so that the minimum intensity value had an intensity of zero and the maximum had an intensity of 1). The MAP2 pixel intensities were enhanced by a tophat Tubeness filter with a scale of 2 sigma of the Gaussian. Once the lower and upper bounds of the three-class Otsu thresholding method were set for the MAP2 pixel detection, the resulting images were converted into segmented objects. The SYNAPSIN1 pixels were enhanced using a tophat Speckles filter, thresholded using a two-class Otsu method, processed for SYNAPSIN1 puncta identification within an equivalent diameter range of 1 to 6 pixels, and de-clumped before being converted into segmented objects. A colocalization module then assigned the relationships between the identified SYNAPSIN1 puncta contained within or only partly touching the MAP2 objects. Finally, the area and the number of the synaptic objects were measured and exported to a *.csv spreadsheet. The CellProfiler pipelines are available at: https://github.com/mberryer/ALPAQAS

### Data analysis

#### Quality control

The output CellProfiler *.csv files were imported in Genedata Screener. As the *.csv files were loaded, a parameterized parser conducted quality control. The mean and standard deviation of the area covered by MAP2 pixels were calculated across all fields in an inter-plate basis. Fields in which the area covered by MAP2 pixels was outside the range of the inter-plate mean ± standard deviation values were removed. Wells containing fewer than 5 fields were also excluded. The area and the number of the synaptic objects of the remaining fields were averaged and condensed to well-level before being imported into Genedata Screener to perform relevant calculations.

#### Synaptic measurements

We analyzed the area occupied by MAP2 positive neurites and the number, area and density of SYNAPSIN1 puncta localized on MAP2 positive neurites (respectively AreaOccupied and Count of MAP2PositiveNeurites and SYNAPSIN1PunctaOnMAP2PositiveNeurites in the ALPAQAS pipeline). On a well-basis, density was defined as the number of SYNAPSIN1 positive puncta colocalized on MAP2 positive pixels divided by the area covered by the MAP2 positive pixels. The mean values for number and density were calculated for each well and expressed as a percentage of the intra-plate DMSO control treated wells. Computed on an intra-plate basis, the Z-score for the value X was defined as the difference between X and the mean value of all DMSO treated wells, divided by the standard deviation value of all DMSO treated wells.

### Statistical analyses

All experiments were performed in three biological replicates and three technical replicates, except the small molecule validation experiments were performed in two biological replicates and eight technical replicates and the siRNA experiment was performed in one biological replicate and six technical replicates. Biological replicates refer to independent batches of cells / differentiations, and technical replicates refer to independent wells. For the dose-response, the small molecule validation, the hN monoculture experiments and the immunoblot quantification, significance was assessed using One-way ANOVA with Dunnett’s multiple comparisons test; *p<0.05, **p<0.01, ***p<0.001. In the hN + hpA co-culture versus hN monoculture experiment, the quantification of the density and the area of the individual SYNAPSIN1 puncta and the area covered by MAP2 positive neurites, the significance was determined by a Kolmogorov-Smirnov unpaired t-test, ***p<0.001. For mRNA-seq analyses, we used an adjusted p-value cutoff of 0.05 and a log_2_fold change cutoff of +/- 1. P values (or adjusted P values where relevant) ≤ 0.05 were considered statistically significant.

### Data, resource and code availability

All data is available in the manuscript or the supplemental materials. The analysis codes generated in this study are available from GitHub at: https://github.com/mberryer/ALPAQAS. Engineered cell lines are available upon request and following appropriate institutional guidelines for their use and distribution.

## ACKNOWLEDGEMENTS

We thank members of the Barrett and Rubin labs for insightful discussions and critical reading of the manuscript. We appreciate the support from Ralda Nehme for her useful inputs on the neuronal induction protocol, Maria Alimova and Patrick Byrne at CDoT for their high-throughput imaging expertise, Sam Bryant and Rachel Fox at the Stanley Center for their analytical insights, Jessica Moffitt for technical support, and Francesca Rapino and Maura Charlton for their wisdom and guidance throughout the project. We also thank Ajamete Kaykas and Katie Worringer for the Ngn2 AAVS1 targeting construct. We appreciate the analytical support from Zach Herbert at the Dana-Farber Cancer Institute Molecular Biology Core Facilities. This work was supported by a Broad*next*10 grant to L.E.B. and L.L.R. as well as support from the Stanley Center for Psychiatric Research. K.W.K., B.A.C. and A.E.C. were supported in part by a grant from the National Institutes of Health NIH R35 GM122547 to A.E.C.

## ETHICS STATEMENT

All studies using hPSCs followed institutional IRB and ESCRO guidelines approved by Harvard University.

## AUTHOR CONTRIBUTIONS

L.E.B. and M.H.B. conceived the project and wrote the manuscript, M.H.B. performed stem cell, neuron and astrocyte cultures, immunostaining, development, optimization and performance of the assays and analyses. S.G.S., D.L. and A.M. generated TALEN edited cell lines, G.R., K.P. and L.L.R provided strategic and technical assistance with screening experiments, D.T. and A.N. performed protein expression analyses, M.H.B., K.W.K., B.A.C. and A.E.C. developed CellProfiler pipelines. L.E.B. and L.L.R. supervised the study and secured funding. All authors discussed results and edited the manuscript.

## SUPPLEMENTAL INFORMATION

**Supplemental Fig. 1:**
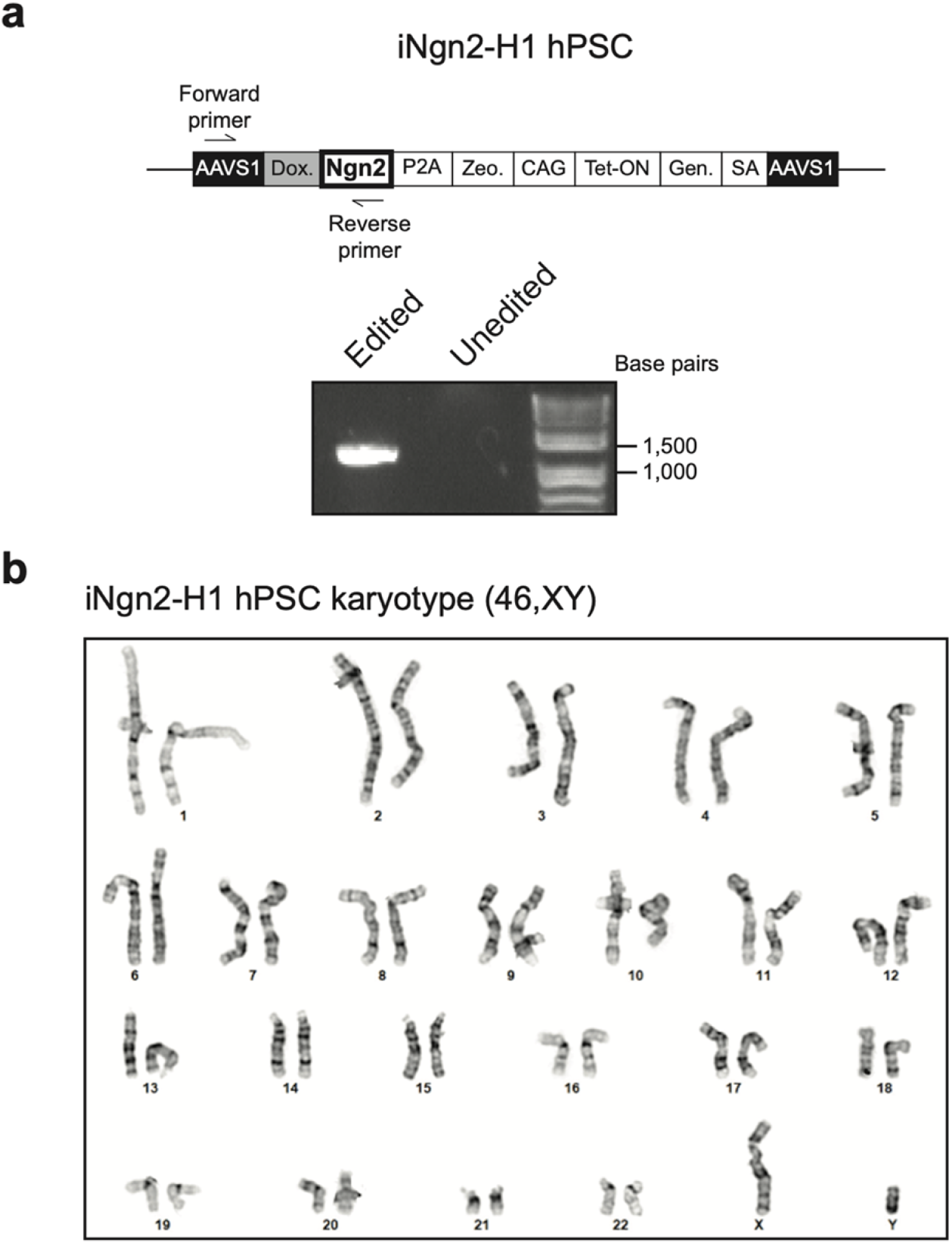
Validation of stably integrated iNgn2 cassette in hPSCs. **a** Design of the PCR genotyping strategy for validation of stable integration of the iNgn2 cassette in the AAVS1 safe harbor locus of the H1 genome. **b** Cytogenetic analysis performed by Cell Line Genetics on iNgn2-hPSCs demonstrating an apparently normal karyotype 46,XY.

**Supplemental Fig. 2:**
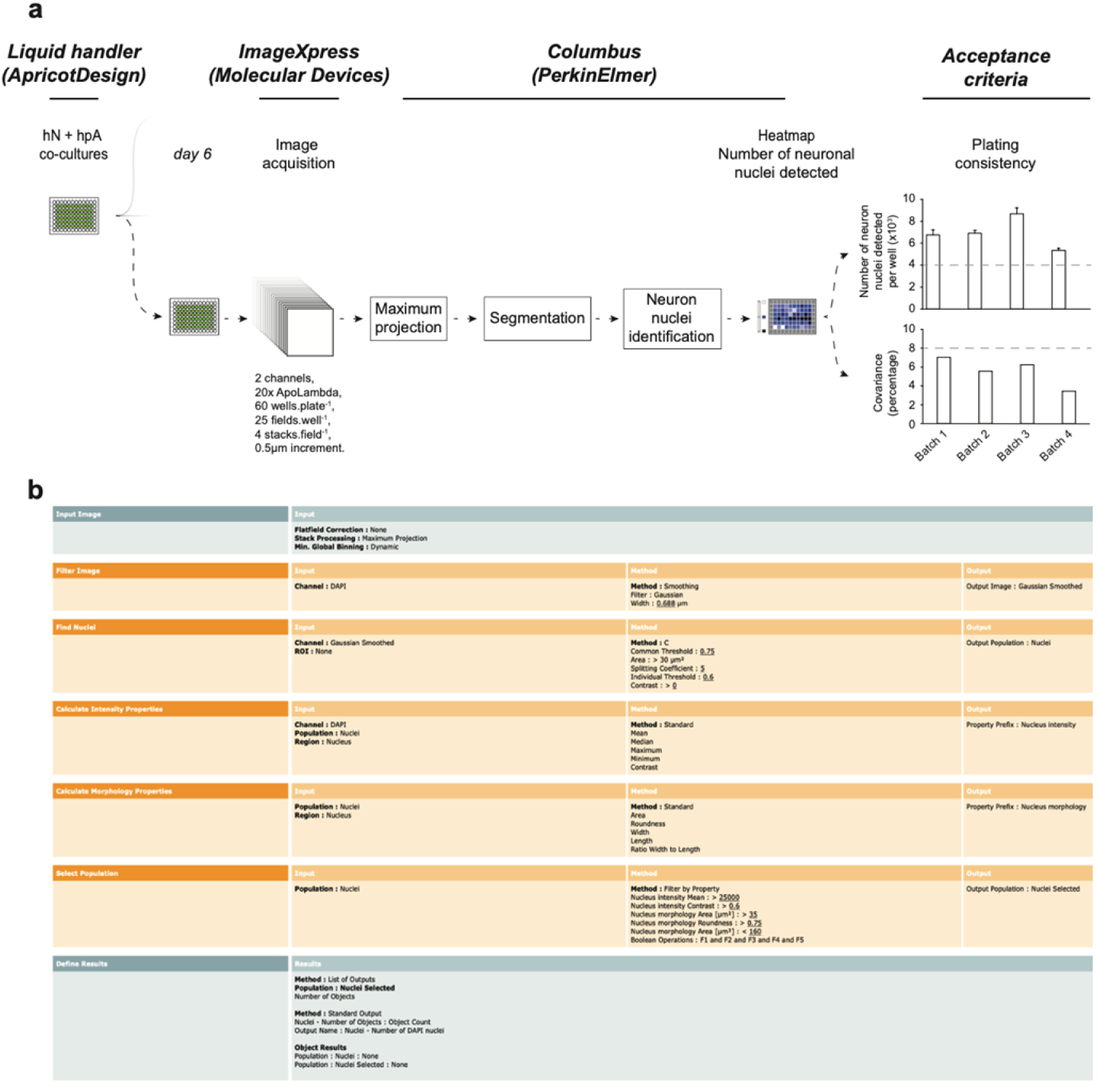
Columbus script to evaluate acceptance criteria for each batch of cultures. **a** One day after seeding cells, a random plate among the batch is selected, stained against MAP2 and counterstained with DAPI. Images are acquired through ImageXpress (Molecular Devices) and uploaded to Columbus (PerkinElmer). DAPI projected images are filtered and the hN nuclei identified based on the intensity, morphology, contrast, area, roundness and MAP2 positive soma. hN nuclei are then counted and the intra-plate covariance is calculated. Acceptance criteria for inclusion of each batch of human co-cultures are set above 4000 hN nuclei detected, and a covariance below 8% as depicted by the grey dashed lines (also shown in Fig. 1). **b** Columbus script for assessment of the number of neuronal DAPI positive nuclei.

**Supplemental Fig. 3:**
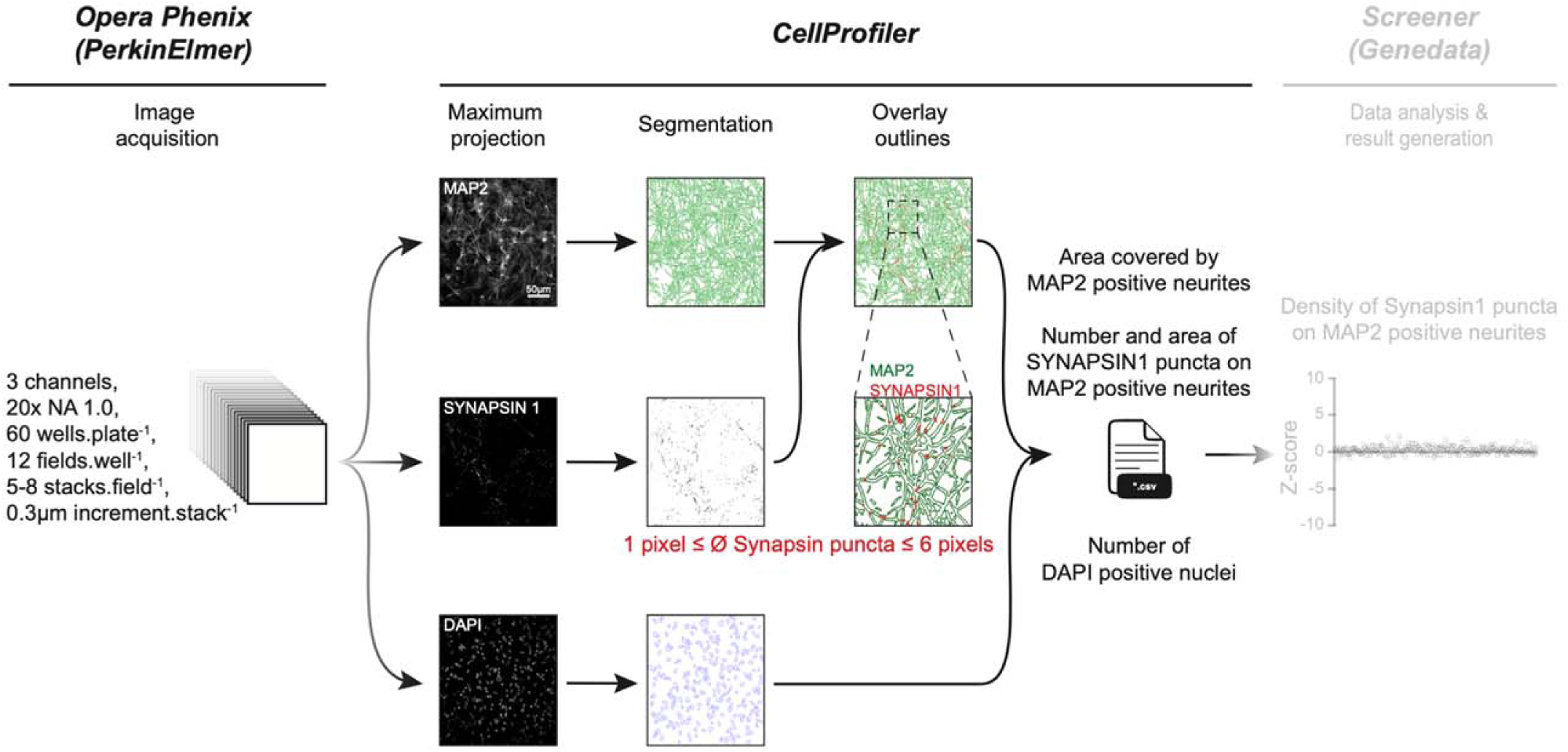
Overview of Automated layered profiling and quantitative analysis of SYNAPSIN1 puncta on MAP2 neurites (ALPAQAS). For each acquired field, five to eight stacks of MAP2, SYNAPSIN1 and DAPI channels are projected in a single plane before segmentation. A colocalization module then assigns the relationship between the identified SYNAPSIN1 puncta contained within or partly touching the MAP2 neurite objects. The area covered by MAP2 neurites and the number of SYNAPSIN1 puncta on MAP2 neurites from each field are then exported to a *.csv spreadsheet.

**Supplemental Fig. 4:**
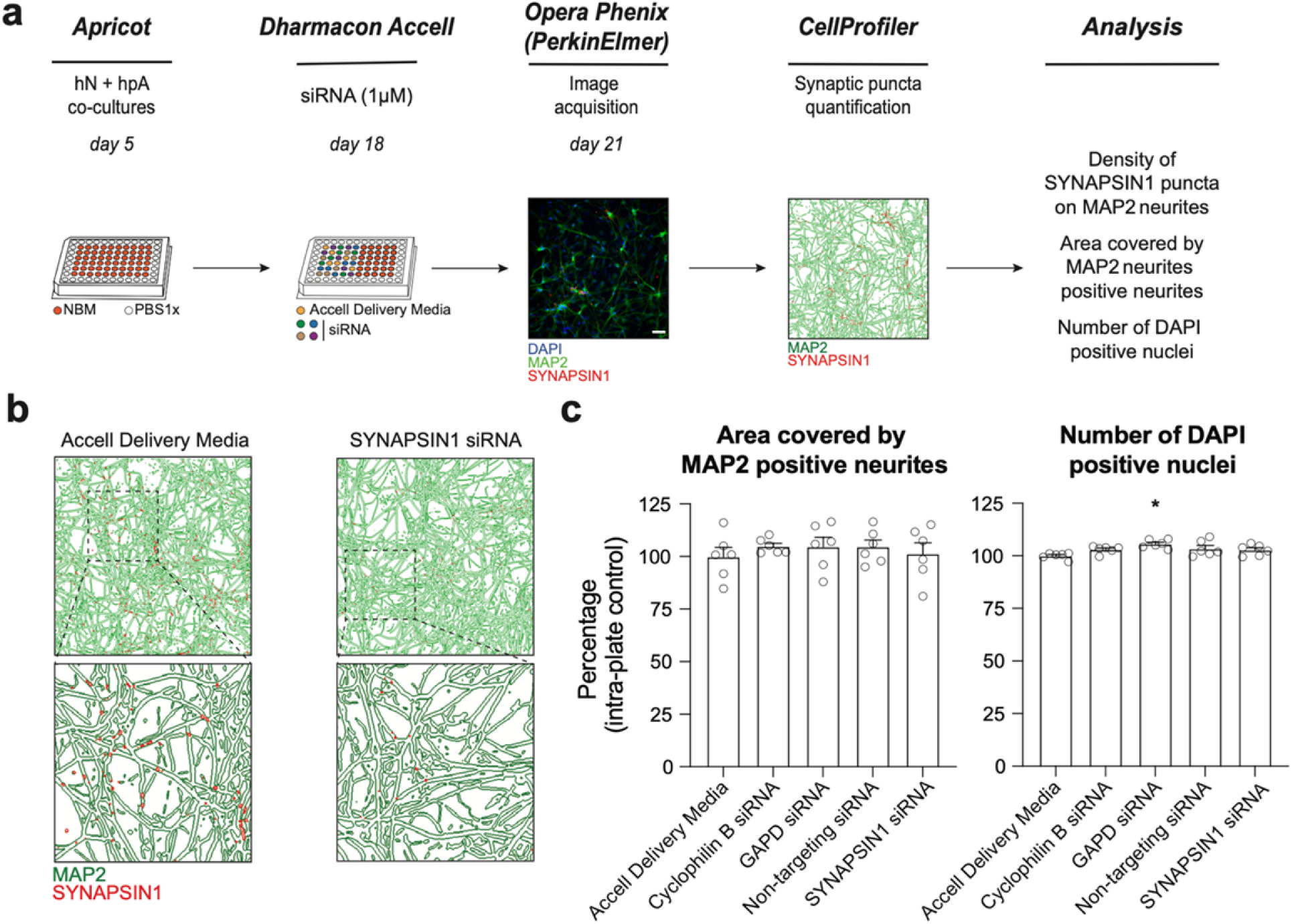
ALPAQAS detects SYNAPSIN1 knock-down. **a** Schematic of siRNA delivery. hN + hpA co-cultures were untreated or incubated on day 18 with SYNAPSIN1 or control siRNA (1µM) for 72 hours, stained and processed through the synaptic assay. **b** Examples of output from CellProfiler showing overlay of SYNAPSIN1 puncta (red) on MAP2 expressing neurites (green) from untreated and SYNAPSIN1 siRNA treated conditions. **c** Histograms showing the MAP2 positive neurite coverage and the number of DAPI positive nuclei after incubation with SYNAPSIN1 siRNA versus control conditions. n = 6 wells for each condition; *p<0.05, ANOVA with Dunnett’s post hoc test.

**Supplemental Fig. 5:**
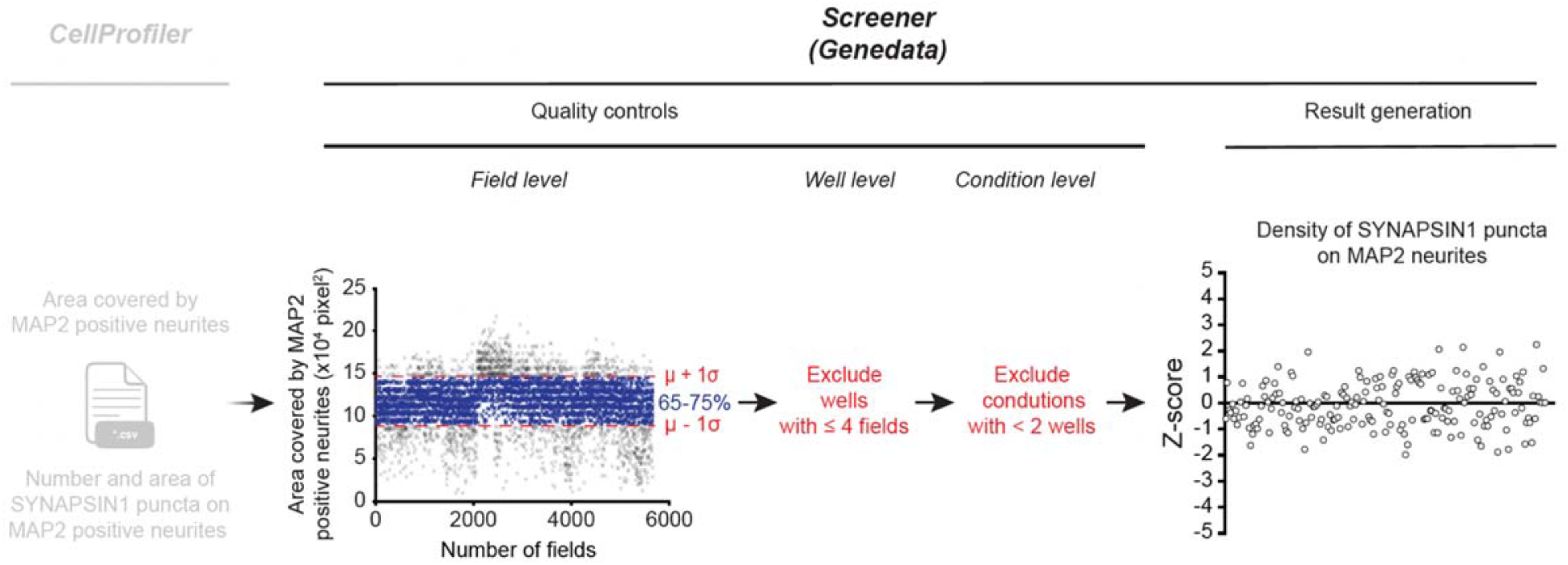
Workflow supported by Genedata. CellProfiler *.csv files (field level) are uploaded with a Genedata parser and filtered through three quality control steps: (*i*) for each individual field (cross) the area covered by MAP2 must fall within ± one standard deviation (σ) to the intra-batch mean (µ) MAP2 coverage (blue crosses represent accepted fields shown within red dashed lines; grey crosses indicated fields that are rejected), (*ii*) exclusion of the well if ≤ 4 fields remain following the first quality control step (iii) exclusion of conditions with one well remaining following the well-level quality control. Data are then imported into Genedata Screener and pattern correction algorithms are leveraged for calculation of Z-score values of the density of SYNAPSIN1 puncta on MAP2 neurites for each condition tested.

**Supplemental Fig. 6:**
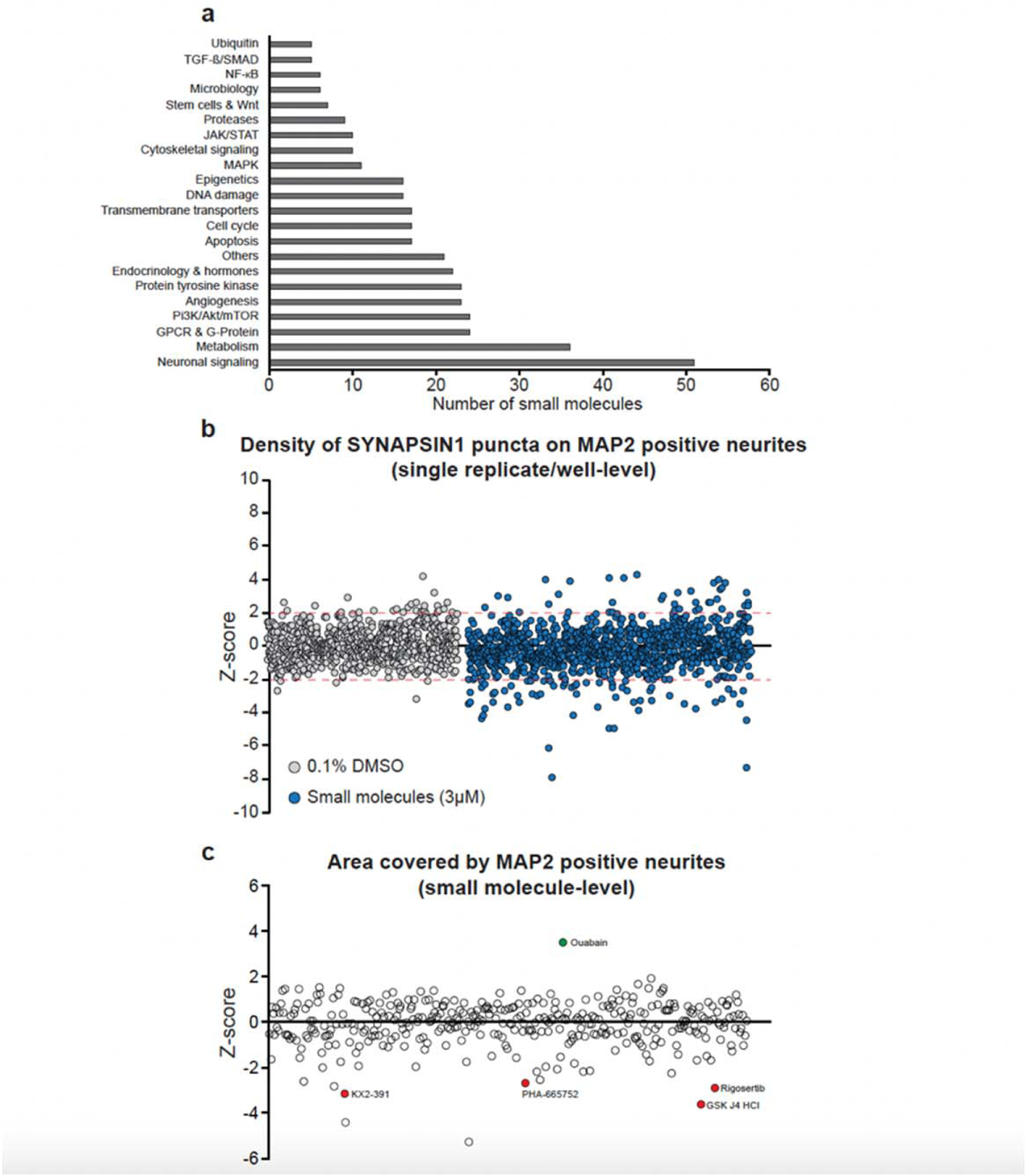
Small molecules used in the primary screen. **a** Number of small molecules per pathway used in the primary screen. **b**, **c** Small molecules were screened at 3µM in triplicate for 72 hours. 13 small molecules did not meet quality control requirements. 92.94% of the control and treated wells had a Z-score for synaptic density between -2 and +2 as depicted by the red dashed lines (**b**). Values are presented as Z-score of the density of SYNAPSIN1 puncta on MAP2 positive neurites (**b**, each circle represents either a DMSO or a small molecule treated well) or the area covered by MAP2 expressing neurites (**c**, each circle represent the aggregate value of the triplicate). Small molecules decreasing the area covered by MAP2 positive neurites are indicated by circles filled in red, and one small molecule increasing the area covered by MAP2 positive neurites is indicated by circle filled in green.

**Supplemental Fig. 7:**
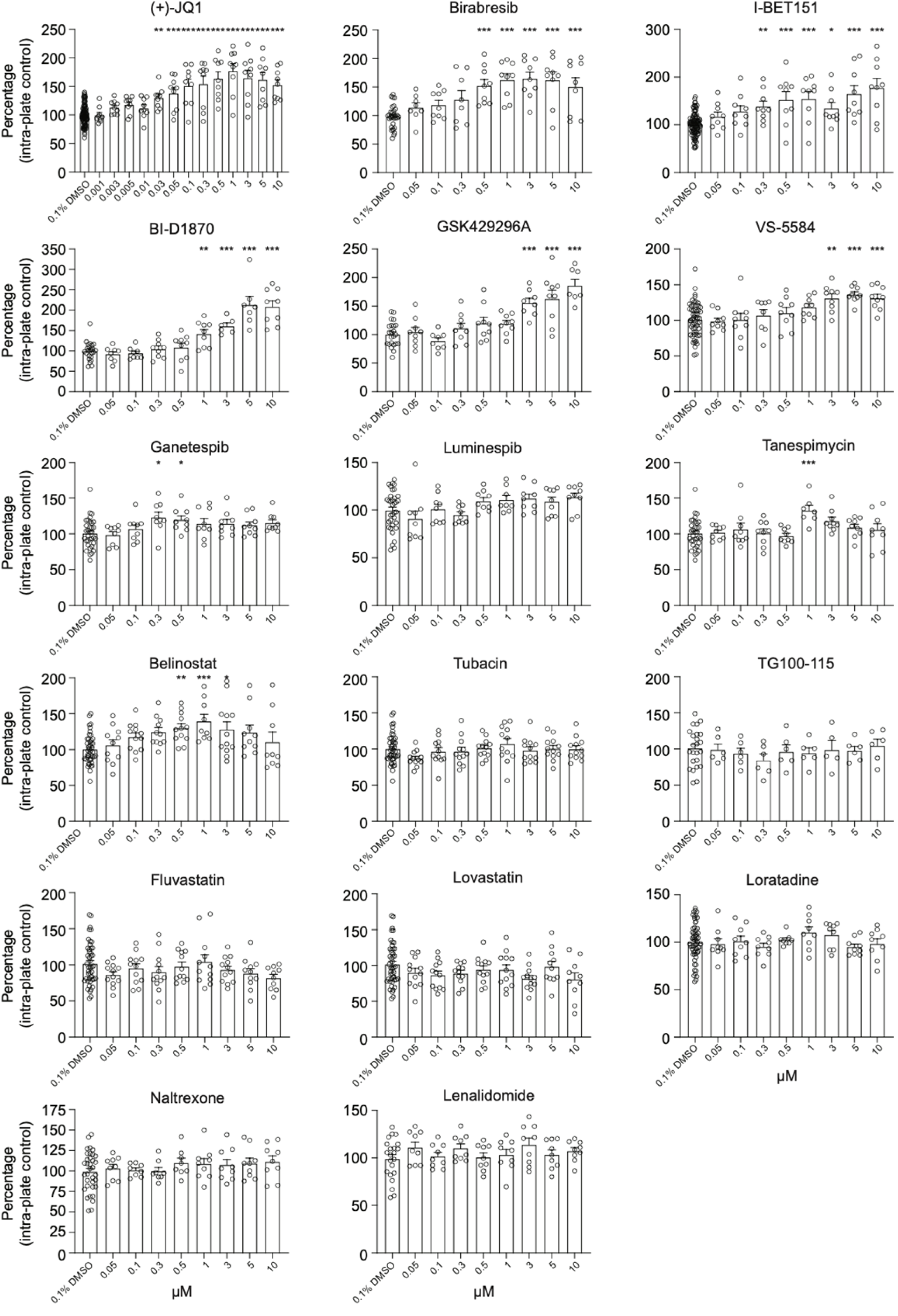
Dose-response histograms for the seventeen selected small molecules on human presynaptic density. Human co-cultures were treated on day 18 with the seventeen selected small molecules at various concentration for 72 hours, stained and processed through the synaptic assay. Histograms showing the density of SYNAPSIN1 puncta on MAP2 expressing neurites. Data are quantified by percentage of intraplate control (0.1% DMSO) represented as mean values ± SEM, n = 3 biological replicates, n = 3 technical replicates. *p<0.05, **p<0.01, ***p<0.001; One-way ANOVA with Dunnett’s multiple comparisons test.

**Supplemental Fig. 8:**
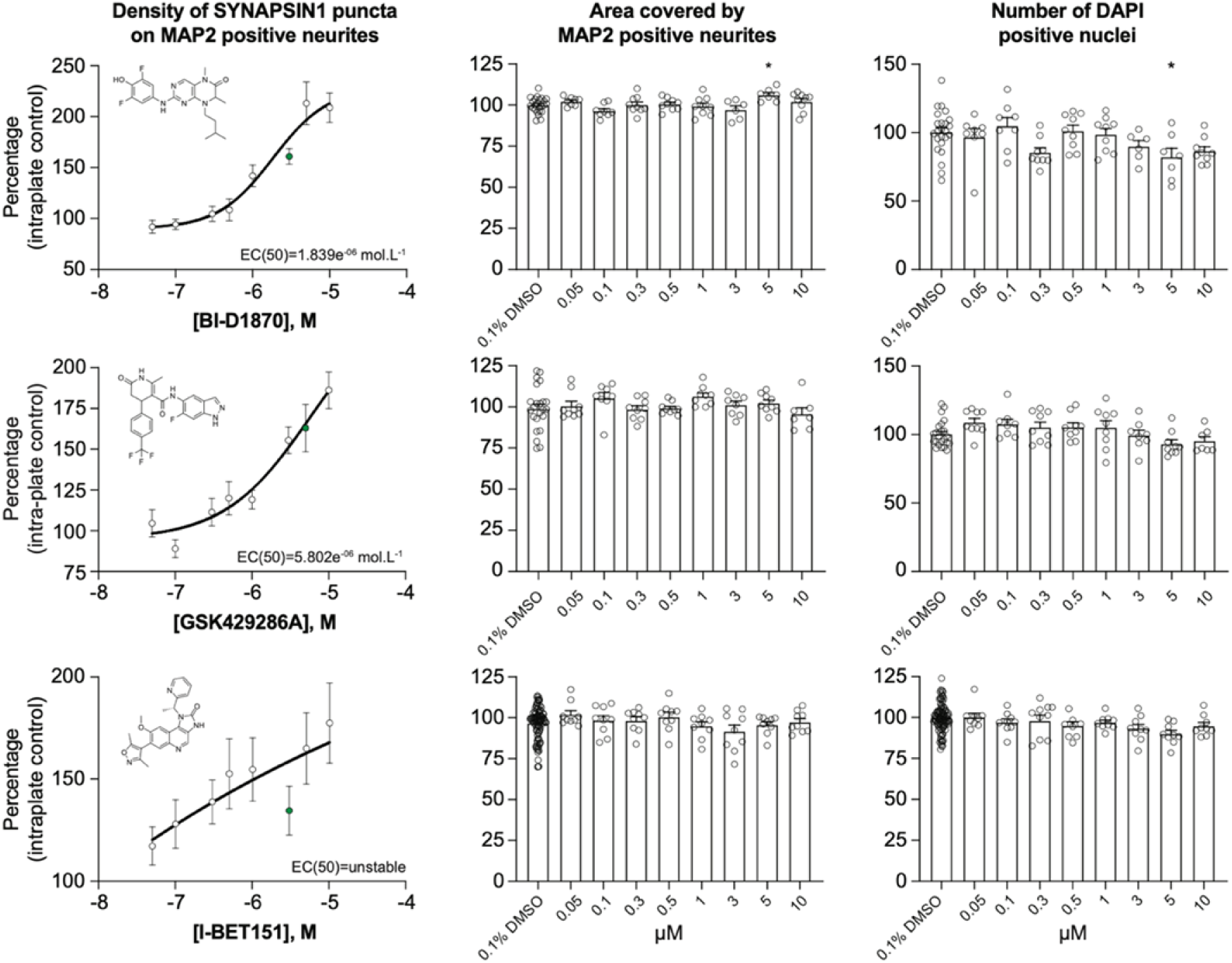
BI-D1870, GSK429286A and I-BET151 increase presynaptic density. Human co-cultures were treated on day 18 with the three selected small molecules at various concentration for 72 hours, stained and processed through the synaptic assay. Concentration responses for BI-D1870, GSK429286A and I-BET151 increasing human presynaptic density (*left*); Green circles indicate the final selected concentration. The impact on the area covered by the MAP2 positive neurites (*middle*) and on the toxicity (number of DAPI positive nuclei; *right*) are also shown. EC(50) and curves were calculated and drawn by PRISM software. Data are quantified by percentage of intraplate control (0.1% DMSO) represented as mean values ± SEM, n = 3 biological replicates, n = 3 technical replicates. *p<0.05; One-way ANOVA with Dunnett’s multiple comparisons test.

**Supplemental Fig. 9:**
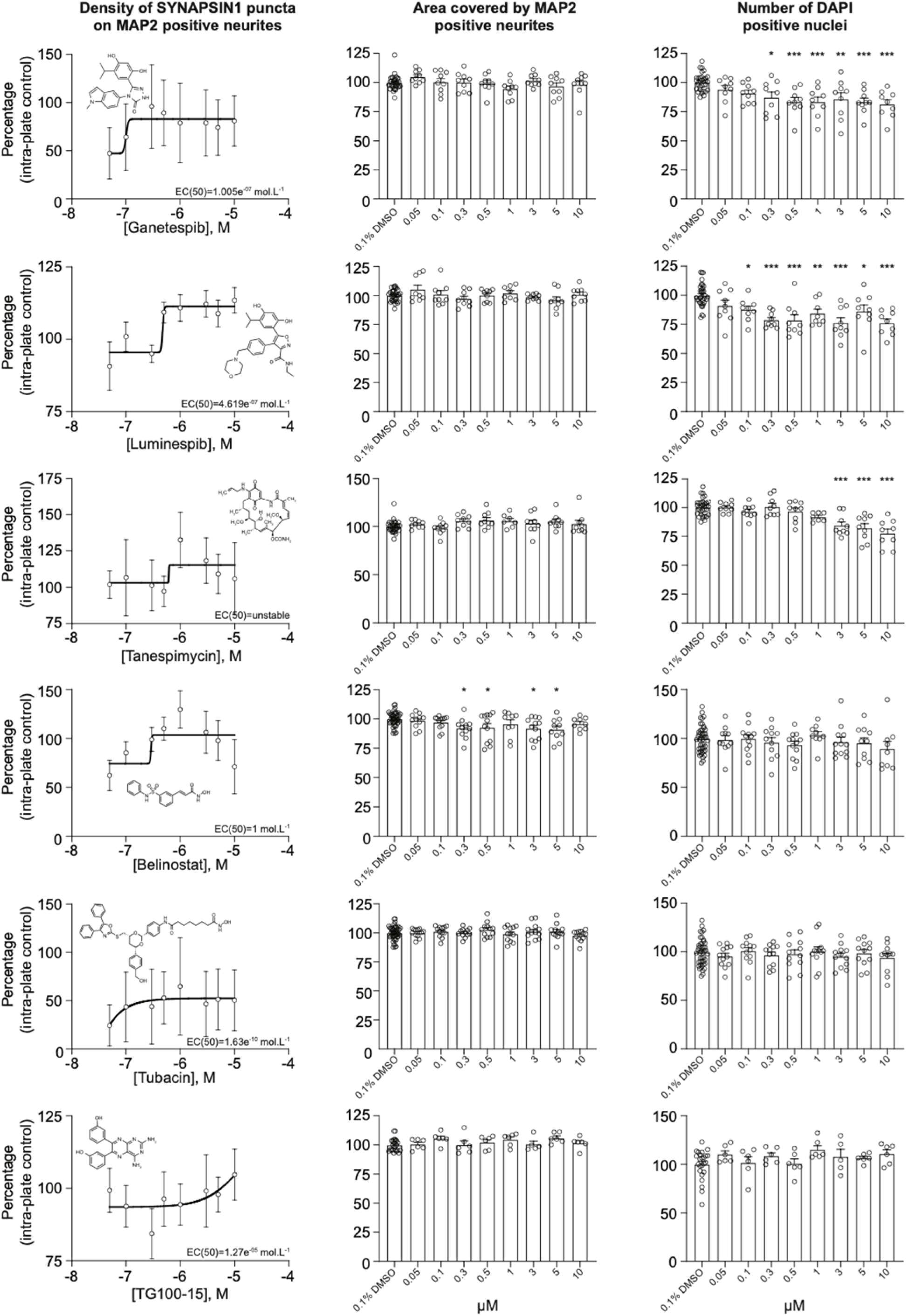
HSP90, HDAC and PI3K inhibitors do not increase human presynaptic density in our assay. Human co-cultures were treated on day 18 with 0.1% DMSO or Ganetespib, Luminespib, Tanespimycin, Belinostat, Tubacin and TG100-15 at various concentration for 72 hours, stained and processed through the synaptic assay. Concentration responses and histograms showing the area covered by MAP2 expressing neurites and the number of DAPI positive nuclei. Each circle represents the value of one replicate. Data are quantified by percentage of intraplate control (0.1% DMSO) represented as mean values ± SEM, n = 3 biological replicates, n = 3 technical replicates. *p<0.05, **p<0.01, ***p<0.001; One-way ANOVA with Dunnett’s multiple comparisons test.

**Supplemental Fig. 10:**
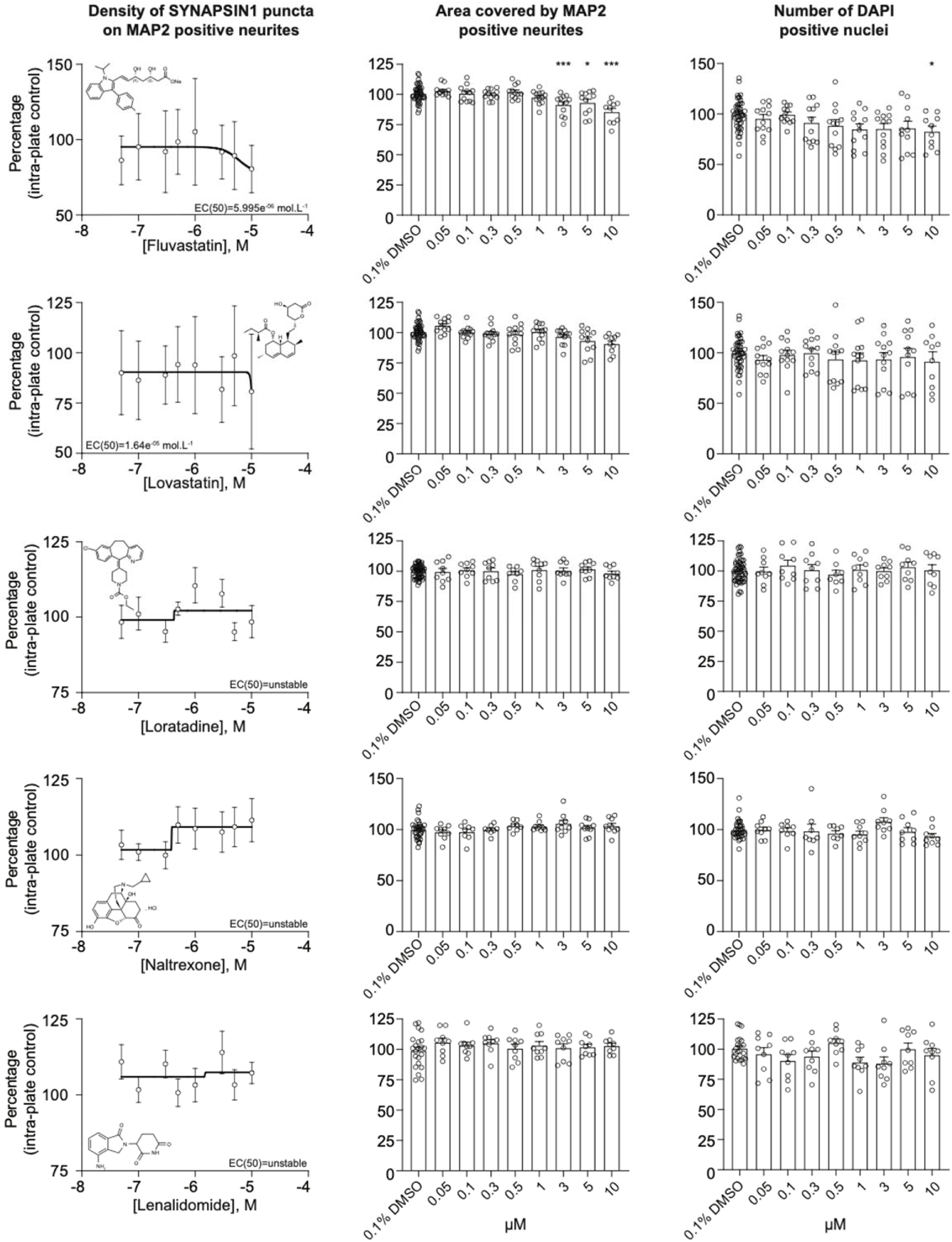
Statins, histamine H1 and opioid receptor antagonists and ubiquitin E3 ligand do not increase presynaptic density in our assay. Human co-cultures were treated on day 18 with 0.1% DMSO or Fluvastatin, Lovastatin, Loratadine, Naltrexone and Lenalidomide at various concentration for 72 hours, stained and processed through the synaptic assay. Concentration responses and histograms showing the area covered by MAP2 expressing neurites and the number of DAPI positive nuclei. Each circle represents the value of one replicate. Data are quantified by percentage of intraplate control (0.1% DMSO) represented as mean values ± SEM, n = 3 biological replicates, n = 3 technical replicates. *p<0.05, ***p<0.001; One-way ANOVA with Dunnett’s multiple comparisons test.

**Supplemental Fig. 11:**
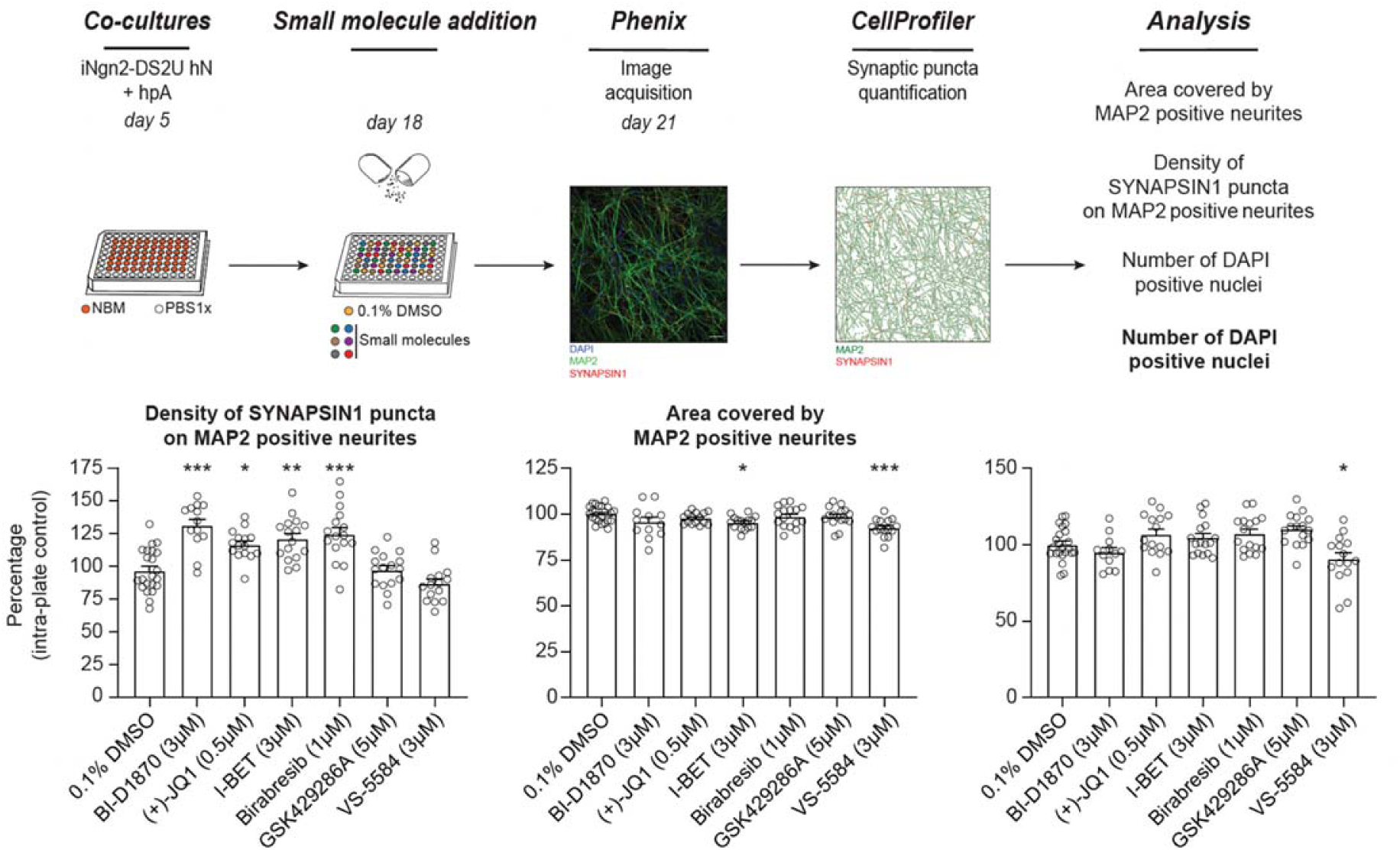
The effect of BET inhibition on presynaptic density is reproducible across cell lines. hNs were generated using an independent cell line (iNgn2-DS2U human iPSCs), co-cultured with hpAs, treated on day 18 with 0.1% DMSO or selected small molecules at the optimal concentrations for 72 hours, stained and processed through the synaptic assay. Histograms show the density of SYNAPSIN1 puncta on MAP2 positive neurons (*left*), the area covered by MAP2 expressing neurites (*middle*) and the number of DAPI positive nuclei (*right*). Data are quantified by percentage of intraplate control (0.1% DMSO) represented as mean values ± SEM, n = 2 biological replicates, n = 8 technical replicates. *p<0.05, **p<0.01, ***p<0.001; One-way ANOVA with Dunnett’s multiple comparisons test.

**Supplemental Fig. 12:**
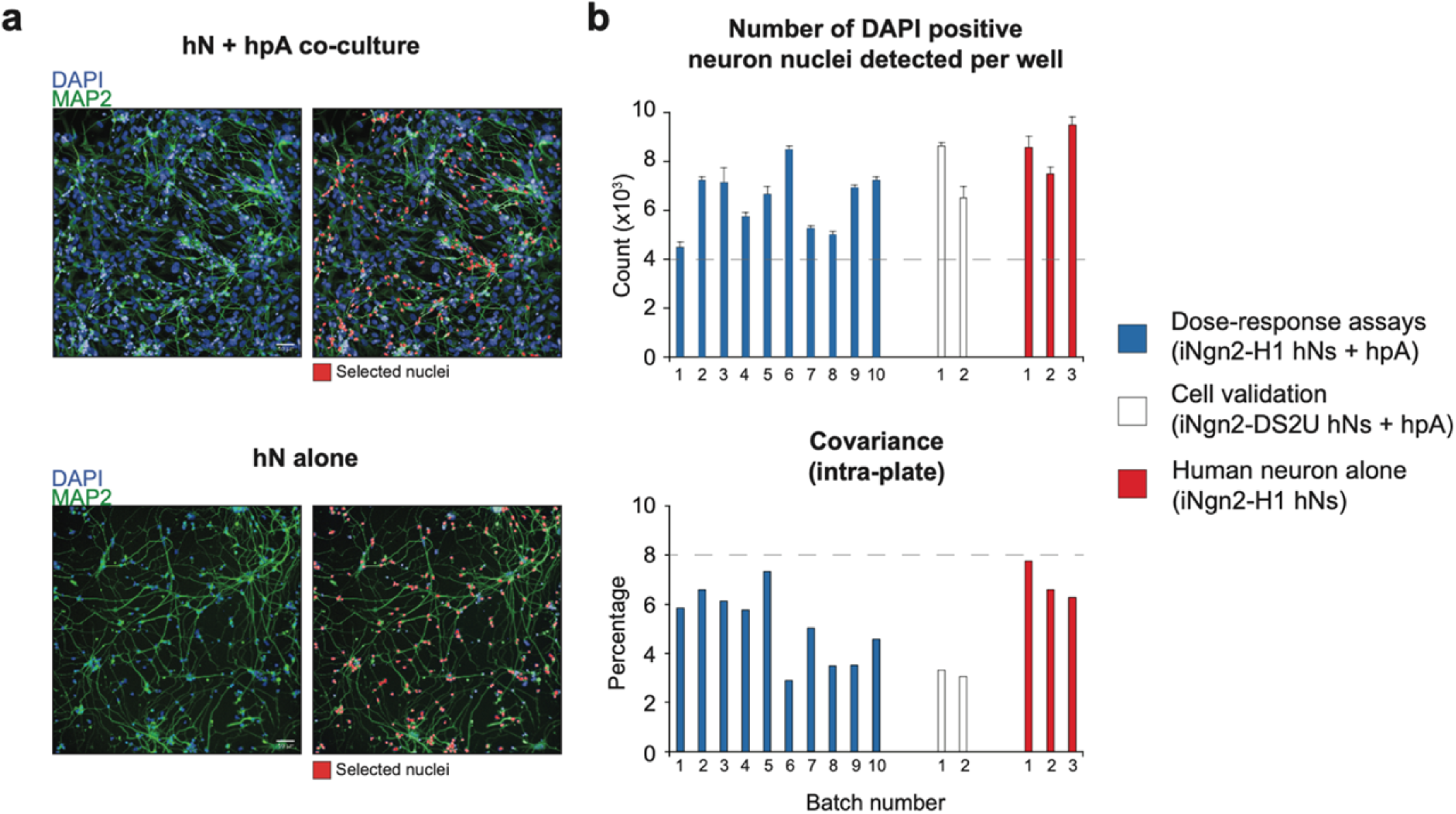
Acceptance criteria of all batches. **a** Representative image of hN + hpA co-culture and hN alone culture fixed one day after plating, stained for MAP2 and counterstained with DAPI. In red, hN DAPI positive nuclei selected through the Columbus script. Scale = 50µm. **b** Number of DAPI positive hN nuclei detected per well and intra-plate covariance for each batch used for the dose-response assays, the small molecule validation, and the hN alone experiments. Acceptance criteria for inclusion of each batch of human co-cultures are set above 4000 hN nuclei detected, and a covariance below 8% as depicted by the grey dashed lines.

**Supplemental Fig. 13:**
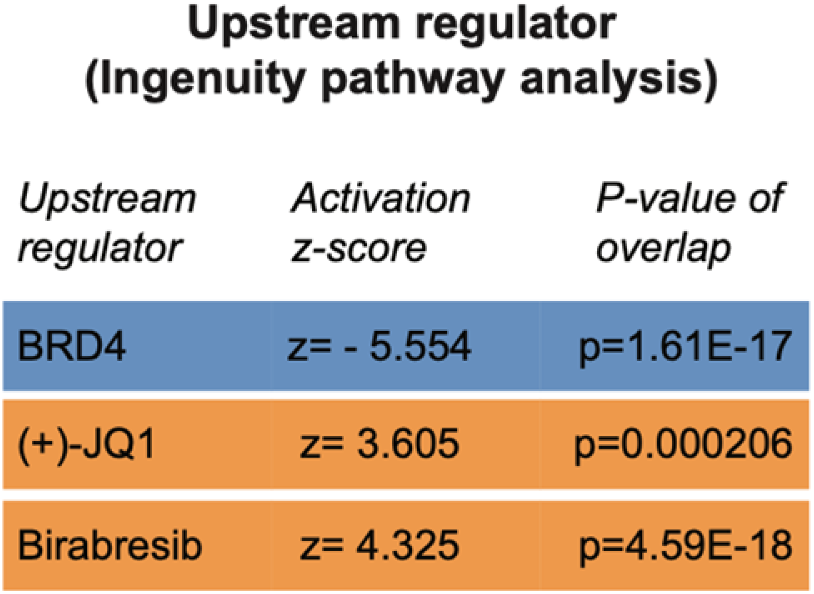
BET proteins and inhibitors are identified as upstream regulators of the DEGs. Significant DEGs shared between (+)-JQ1 and Birabresib treatment conditions were uploaded in Ingenuity Pathway Analysis, and the activation z-scores and p-values of the overlap were quantified. Blue indicates a negative activation z-score (BRD4) and orange indicates a positive activation z-score ((+)-JQ1 and Birabresib).

**Supplemental Table 1: List of the 376 small molecules used in the primary screen.**

**Supplemental Table 2: Transcriptional analyses after (+)-JQ1 inhibition.**

**Supplemental Table 3: Transcriptional analyses after Birabresib inhibition.**

